# A toolbox for efficient proximity-dependent biotinylation in zebrafish embryos

**DOI:** 10.1101/2021.05.16.444353

**Authors:** Shimon M. Rosenthal, Tvisha Misra, Hala Abdouni, Tess C. Branon, Alice Y. Ting, Ian C. Scott, Anne-Claude Gingras

## Abstract

Understanding how proteins are organized in compartments is essential to elucidating their function. While proximity-dependent approaches such as BioID have enabled a massive increase in information about organelles, protein complexes and other structures in cell culture, to date there have been only a few studies in living vertebrates. Here, we adapted proximity labeling for protein discovery *in vivo* in the vertebrate model organism, zebrafish. Using lamin A (LMNA) as bait and green fluorescent protein (GFP) as a negative control, we developed, optimized, and benchmarked *in vivo* TurboID and miniTurbo labeling in early zebrafish embryos. We developed both an mRNA injection protocol and a transgenic system in which transgene expression is controlled by a heat shock promoter. In both cases, biotin is provided directly in the egg water, and we demonstrate that 12 hours of labeling are sufficient for biotinylation of prey proteins, which should permit time-resolved analysis of development. After statistical scoring, we found that the proximal partners of LMNA detected in each system were enriched for nuclear envelope and nuclear membrane proteins, and included many orthologs of human proteins identified as proximity partners of lamin A in mammalian cell culture. The tools and protocols developed here will allow zebrafish researchers to complement genetic tools with powerful proteomics approaches.

## Introduction

Probing the organization of proteins into complexes, organelles and other structures in the context of living organisms is critical to understanding the context-dependent nature of protein function. While unicellular model organisms, particularly the yeast *Saccharomyces cerevisiae*, have been well characterized, the organization of the proteome in the living cells of multicellular organisms including vertebrates is still largely uncharted. The vertebrate model organism *Danio rerio* (zebrafish) is a popular model to interrogate vertebrate development. While traditionally used for developmental studies due to its high fecundity, external fertilization, rapid development, and optical clarity during early development (1), recently the use of zebrafish for disease modeling and drug screening has increased. This has spurred a coincident increase in genetic tools adapted or developed for zebrafish use. However, the adaptation and development of tools for proteomic studies in zebrafish has lagged behind.

In zebrafish, there are two main methods by which exogenous protein expression can be driven, either by the injection of mRNA or through the generation of transgenic lines (reviewed in (2)). Each approach has its strengths and limitations. Injection of *in vitro* transcribed mRNA is usually performed at the single cell stage to achieve expression throughout the developing embryo. The advantage is that there is no lag time in performing an experiment as there would be when generating a transgenic animal. This method is most appropriate when early development (<72 hours post fertilization (hpf)) is being studied, as consistent expression levels are difficult to maintain for much longer due to mRNA and protein degradation (3). A potential drawback, particularly in cases where many embryos are required, is that each embryo needs to be injected individually, a time-consuming task and one potentially leading to variability of protein expression levels. Finally, tissue specific expression is not possible via mRNA injection. Transgenics are ideal for later time points and should afford versatility in both expression levels and site of expression when employing different promoters, e.g. stage specific or tissue specific (reviewed in (4)). Disadvantages of transgenics are the relatively lengthy generation time of 2.5–4 months, and the resultant overall difficulty in scaling up the number of genes profiled by the approach.

Studies of proteome organization have typically employed biochemical fractionation and affinity purification, both coupled to mass spectrometry. These methods are ideal for studying stable interactions after tissues and cells have been lysed, but they require that the protein complexes or organelles stay intact throughout the purification (5). Additionally, the cellular context is disrupted upon lysis, which leads to loss of information as well as the risk for post-lysis artefacts.

The introduction of proximity-dependent biotinylation techniques in 2012 has offered new possibilities for the detection of protein-protein interactions, including weaker interactions, and the definition of the composition of both membrane-bound and membraneless organelles (reviewed in (6)). Proximity-dependent biotinylation approaches include biotin ligase-based methods such BioID (7) and peroxidase-based methods like APEX (8) that permit the covalent labeling of preys proximal to a protein of interest in the context of living cells. Biotin ligase-based methods take advantage of a mutated biotin ligase, either an *E. coli* BirA harboring a single point mutation [R118G, referred to as BirA* in the original BioID study (7)], molecularly evolved derivatives called miniTurbo and TurboID – see below – or mutated biotin ligases from other species such as BioID2 from *Aquifex aeolicus* or BASU from *Bacillus subtilis* (11). These mutated enzymes are capable of activating biotin to the reactive intermediate biotinyl-5’-AMP, but instead of completing the transfer of biotin to a biotin-acceptor peptide, they release it from the active site where it can react with primary amines in the vicinity, resulting in their covalent biotinylation (reviewed in (12)). Fusion of the mutated biotin ligase to a protein of interest (the “bait”), expression of the fusion in a cell of interest, and addition of exogenous biotin therefore lead to the labeling of proteins within a ∼10 nm radius from the bait (13). The labeled proteins (preys) include a mixture of direct and indirect interactors (often including weak or cycling interacting partners) as well as proteins that reside in the same locale as the bait (collectively referred to as “proximal interactors”). Because the biotin is covalently bound, harsh lysis, solubilization, and washes can be performed, allowing for the recovery of previously inaccessible proximal interactors.

Proximity-dependent biotinylation has been employed by numerous groups to investigate the proximal interactomes of proteins, complexes and organelles in cells grown in culture. While BioID with the original BirA* enzyme (7) or the *Aquifex aeolicus* biotin ligase BioID2 (10) have been performed in unicellular and multicellular organisms such as *Dictyostelium discoideum* (14), *Plasmodium berghei* (15)*, Toxoplasma gondii* (16), *Trypanosoma brucei* (17), *Saccharomyces cerevisiae* (18), and *Nicotiana benthamiana* (19), there have been relatively few reports of their use in vertebrates, most of them in mice (20–25). Pronobis *et al*. used BioID2 to study heart regeneration in transgenic zebrafish following repeated injections of biotin intraperitoneally over 3 days (26), demonstrating that zebrafish is an amenable model for proximity-dependent biotinylation.

Major hurdles to the application of BioID with first generation enzymes in larger animal models relate both to the relatively poor activity of these enzymes (forcing long labeling times as in the zebrafish BioID2 experiment (26)), and the need to deliver supplemental biotin to the appropriate tissues or organs. The generation of molecularly evolved variants of the *E. coli* BirA enzyme, called TurboID and miniTurbo, that display 15 to 30 times the labeling efficiency of BirA* has been a major breakthrough in enabling more efficient proximity-dependent biotinylation in different *in vivo* models, including *C. elegans* (9), *D. melanogaster* (9, 27), *A. thaliana* (28), *N. benthamiana* (28, 29), *S. pombe* (30), and *M. musculus* (31). Using transgenics in zebrafish, Xiong *et al*. recently demonstrated the applicability TurboID through the fusion of a GFP nanobody to TurboID: crosses to transgenic lines expressing fusions of proteins of interest to the Clover green fluorescent protein enabled labeling and recovery of proximal interactors to cavin proteins (32). Yet, proximity-dependent biotinylation approaches are still in their infancy, and more thorough optimization and benchmarking of the approaches is needed; it has also not been clearly determined whether the more rapid and scalable mRNA injection systems lead to high-quality proximal proteomes.

Here, we have adapted proximity-dependent biotinylation with the miniTurbo and TurboID enzymes for use in live zebrafish embryos and demonstrate its versatility as a method that can be applied both via mRNA injection and generation of transgenic animals. To benchmark our method, we demonstrate that we can identify known proximal interactors of the nuclear envelope protein, lamin A. This establishes a toolbox and methodology where any bait can be examined in a rapid fashion in zebrafish embryos via the mRNA injection-based approach, or more slowly but with greater nuance using the transgenic approach. This will be of particular value for proteins whose function is cell- or context-dependent.

## Experimental Procedures

### Generation of zebrafish vector sequences and expression constructs

The TurboID and miniTurbo sequences (9) were optimized for zebrafish expression using the online codon optimization tool from Integrated DNA Technologies (IDT). A zebrafish consensus Kozak sequence shown to generate high expression (33) was added at the start site (GCAAACatgGCG). A flexible linker (GS2) was included downstream of the protein coding sequence followed by a 3x FLAG tag. Finally, a Gateway cloning attR1 site was added at the 3’ end of the sequences. Upstream and downstream restriction sites were also included to allow for restriction cloning. For the humanized version, the constructs were identical except for the use of a generic Kozak sequence (CGCCACCatgGCG).

The above constructs were synthesized and cloned into the pUC57 vector (General Biosystems Inc. Morrisville, NC, USA). A fusion construct was built in the pcDNA5-pDEST vector. Briefly, the synthesized constructs were restriction cloned (NheI/NotI) into the pcDNA5-pDEST vector upstream of Gateway cloning attR2 sites. A Gateway-compatible entry clone for LMNA (Genbank accession number EU832167, encoding human lamin A) was transferred into the pcDNA5-pDEST Gateway destination vector using LR Gateway cloning. This resulted in a mammalian expression vector of LMNA tagged with TurboID or miniTurbo at its N-terminus. Vectors were constructed in the identical fashion for controls using EGFP (Enhanced green fluorescent protein), by subcloning the EGFP cassette from pDONR233-eGFP (itself a derivative of pEGFP-C1, BD Biosciences). All the constructs were validated by restriction digest and DNA sequencing of the subcloned fragments.

The cassettes (TurboID or miniTurbo fused to LMNA or GFP) were amplified by PCR from pcDNA5-pDEST and transferred to vectors compatible with zebrafish expression. First, restriction cloning was used to transfer the cassettes into the pCS2+ vector for mRNA expression (SP6-based) for injection in zebrafish. Briefly, PCR amplified cassettes were digested with ClaI and XbaI and ligated into the pCS2+ vector downstream of a SP6 promoter sequence and upstream of a SV40 polyA cassette. The vectors were validated by restriction digest and DNA sequencing of the subcloned fragments.

Alternatively, the generation of constructs for transgenic generation used the Gateway compatible Tol2kit (34). The Tol2kit is a transposon-based transgenesis method, whereby Gateway cloning is used to insert fragments containing 3’, middle, and 5’ inserts into a destination vector containing flanking Tol2 transposon recognition sites. Upon injection of the vector together with mRNA coding for the Tol2 transposase, the cassette is randomly inserted into the genome (34). The cassettes (TurboID or miniTurbo fused to LMNA or GFP) were amplified by PCR as above and inserted into the Gateway compatible Tol2kit middle entry vector pME-MCS (34) via restriction cloning as above. This vector contains the multiple cloning site derived from the pBluescript II SK+ vector inserted into the pDONR221 vector (34). Tol2 destination vectors were generated using three-fragment Gateway cloning. The Tol2kit destination vector pDestTol2CG2 (34) was used for all constructs as it includes a cardiomyocyte-specific GFP expression sequence, allowing for visual selection of integration positive embryos. A three-fragment Gateway LR reaction was used to generate the final destination vectors. At the 5’ end, the Tol2kit p5E-HSP70l (34) vector was used to donate the 1.5 kb HSP70l promoter fragment (which enables heat shock-inducible expression). This was followed by the fusion protein sequence cassette (TurboID or miniTurbo fused to LMNA or GFP) obtained from the pME-MCS vector. Lastly, an SV40 polyA sequence was obtained from the Tol2kit P3E-polyA (34) vector. All the constructs were validated by restriction digest and DNA sequencing of the subcloned fragments.

### Zebrafish husbandry

Zebrafish were all maintained under the guidance and approval of the Canadian Council on Animal Care and the Hospital for Sick Children Laboratory Animal Services. Embryos were maintained at 28.5°C in embryo medium (egg water). When used, biotin supplementation was performed by directly dissolving biotin powder (BioBasic, cat. BB0078) in heated (∼90℃) egg water followed by cooling and adjustment of the pH to 7. Wild type (WT) or transgenic fish of the strain TL/AB were used for all experiments.

### mRNA injection-based expression of fusion proteins

For expression of fusion proteins via mRNA injection, the pCS2+ vectors containing the fusion constructs were linearized by digestion with NotI. Capped mRNA was generated from the linearized vector using the mMESSAGE mMACHINE SP6 Transcription Kit (ThermoFisher Scientific cat. AM1340) and the transcribed mRNA purified using the MEGAclear Transcription Clean-Up Kit (ThermoFisher Scientific cat. AM1908). The mRNA was quantified by spectrophotometry (NanoDrop 2000) and run on an agarose gel to determine that only a single band was present (to ensure degradation had not occurred). For protein expression, 400 pg of capped mRNA was injected into the yolk of single cell stage wild type embryos. For western blots, a pool of ∼100 deyolked (see below “*Preparation of embryos for protein extraction”*) embryos at 48 hours post fertilization (hpf) was used to generate lysate, with 50 µg of pooled lysate (equivalent to ∼10 embryos) run on the gel per sample. For TurboID pulldowns, ∼1000 and ∼200 deyolked embryos were used for 12 and 48 hpf collections, respectively. In order to properly synchronize the staging of the pool of ∼1000 12 hpf embryos, the injections and collections were staggered. ∼250 embryos were injected per mating tank over ∼15 minutes, after which the injections were repeated with a new mating tank. The injected embryos were incubated at 28.5℃ in biotin-supplemented egg water, and collected 12 hours after their respective injection times. For TurboID, all experiments were performed in biological replicates (duplicates or triplicates as described in the relevant figures), with separate injections performed on different days.

### Generation of transgenic zebrafish lines and protein expression

To generate transgenic zebrafish lines, 30 ng of the final Tol2 destination vectors (purified using the QIAprep Spin Miniprep Kit; Qiagen cat. 27106) were injected into single cell stage zebrafish embryos together with 75 ng of Tol2 transposase mRNA (purified using the MEGAclear™ Transcription Clean-Up Kit; ThermoFisher Scientific cat. AM1908). The injected embryos were screened for positive construct integration based on the presence of GFP expression in the heart at 24–48 hpf (4). GFP-positive embryos were grown to adulthood. To determine if the GFP positive adults had germ-line integration of the constructs, the fish were outcrossed to wild type fish and the progeny screened for GFP expression in the heart. These F1 GFP positive progeny were grown to adulthood. Embryos from incrosses of F1 adults were used for experiments.

Embryos were kept at 28.5℃ until induction of transgene expression by heat shock. At the selected time point for transgene induction (60 hpf), the embryos were transferred to 50 ml conical tubes filled with egg water pre-heated to 38℃ and placed in a 38℃ water bath for one hour (35). Following heat shock, the embryos were returned to plates with 28.5℃ egg water and incubated for the times indicated. Previous studies have found mRNA expression to be induced as early as 15 minutes post heat shock, while protein expression follows at 1.5–4 hours post heat shock. While the stability of the mRNA transcript varies greatly (degrading as soon as three hours post heat shock), protein expression is generally more stable (36–38).

### Preparation of embryos for protein extraction

Following protein expression via mRNA injection or heat shock induction, labelling was allowed to proceed at 28.5℃. Labeling was stopped by washing the embryos in ice-cold egg water. Embryos were dechorionated using pronase in egg water (1 mg/ml; Roche, Cat# 10165921001). Embryos were washed three times with ice-cold egg water and transferred to 1.5 ml microcentrifuge tubes. Embryos were deyolked essentially as described (40). Briefly, embryos were washed twice with ice-cold deyolking buffer (55 mM NaCl, 1.8 mM KCl, 1.25 mM NaHCO_3_). 200 µL deyolking buffer supplemented with 1mM phenylmethylsulfonyl fluoride (PMSF; BioShop Canada, Cat# PMS123) and protease inhibitor cocktail (Sigma-Aldrich, Cat# P8340, 1:100) were added and the yolk disrupted by gentle pipetting using a p200 tip. The yolk was further solubilized by vortexing three times for 2 seconds. The embryos were spun down (300g) and the supernatant removed. Embryos were washed twice with ice-cold wash buffer (110 mM NaCl, 3.5 mM KCl, 2.7 mM CaCl_2_, 10 mM Tris-HCl pH 8.5). Following removal of the supernatant, the embryos were snap frozen in liquid nitrogen, and stored at −80℃.

### Immunoblotting

For western blot analysis of mRNA injected embryos, 50 µg of pooled lysate from ∼100 48 hpf embryos was used, while for transgenic lines, 50 µg of pooled lysate from ∼100 72 hpf embryos was used. Protein extraction was performed as for TurboID affinity purification (see below) with the following modification: the volume employed was 1 µL per embryo for 48 and 72 hpf embryos. A BCA assay (ThermoFisher Scientific, Cat# 23227) was used to quantify protein concentration in the lysate. The lysate was boiled in Laemmli SDS-PAGE sample buffer for five minutes and 50 µg of protein per lane were separated onto 10% SDS-PAGE gels and transferred onto nitrocellulose membranes (GE Healthcare Life Science, Cat# 10600001). Following Ponceau S staining, blots were blocked in 4.5% milk (for anti-BirA probing) or 5% bovine serum albumin (for Streptavidin-Horseradish peroxidase (HRP) probing) in TBS with 0.1% Tween-20 (TBST). Bait proteins were probed using rabbit anti-BirA antibody (Sino Biological, Cat# 11582-RP01) at 1:2000 in block buffer, washed in TBST, and detected with 1:5000 Donkey anti-Rabbit IgG-HRP. Biotinylated proteins were probed using Streptavidin-HRP (GE Healthcare Life Science, Cat# RPN1231) at 1:2000 in blocking buffer. Membranes were developed using ECL™ Western Blotting Detection Reagents (GE Healthcare Life Science, Cat# RPN2109). Quantification of western blots was performed using the Image Studio^TM^ Lite software (Version 5.2, LI-COR Biosciences)

### Live imaging and whole mount immunofluorescence

For imaging GFP expression post heat shock, embryos were subjected to heat shock at 38℃ for one hour. At six hours post heat shock, the embryos were sedated by the addition of tricaine methanesulfonate (0.01 mg/ml) (Sigma-Aldrich, Cat# A5040) and were imaged using a Zeiss Axio ZoomV16 Stereo microscope. Following imaging, the embryos were placed in fresh egg water.

Embryos which were to be used in immunofluorescence experiments were grown in egg water supplemented with 0.003% 1-phenyl 2-thiourea (PTU) (Sigma-Aldrich, Cat# P7629) from 24 hpf to prevent pigmentation. Following induction by heat shock of protein expression at 60 hpf and 12 hours of labeling, 72 hpf embryos were collected for immunofluorescence. Fixing and imaging were performed essentially as in reference (39), with the following modifications. Embryos were washed three times in ice-cold egg water prior to fixation. Fixed embryos were washed twice in PBS + 0.1% Tween-20 (PBSTw) prior to stepwise dehydration using methanol in PBSTw (25%, 50%, 75%, 100% methanol). Following rehydration (performed as dehydration but in reverse), permeabilization was performed with ice-cold acetone at −20℃ for 20 minutes, followed by three 10 minutes washes with PBS + 1% Triton-X100. Embryos were blocked with 10% Normal Goat Serum (NGS; Millipore, Cat# S26-LITER) and 5% Bovine Serum Albumin (BSA) in PBS + 0.4% Triton-X100 (PBSTx) for 1.5 hours at room temperature before being stained for bait protein with mouse anti-FLAG antibody (Monoclonal anti-FLAG M2 antibody; Sigma-Aldrich, Cat# F3165; 1:500) in blocking buffer overnight at 4℃. Embryos were washed six times for 30 minutes in PBSTx. For secondary detection, embryos were stained with goat anti-mouse Alexa Fluor^TM^ 488 (Molecular Probes, ThermoFisher Scientific, A-11032; 1:200), and for visualizing protein biotinylation, with Alexa Fluor^TM^ 594 streptavidin conjugate (Molecular Probes, ThermoFisher Scientific, S21374; 1:500). DAPI (1:1000) was used as a nuclear counterstain. All secondary staining was performed in blocking buffer overnight at 4℃. Embryos were washed six times for 30 minutes in PBSTx and stored at 4℃ in PBSTw. For imaging, embryos were mounted in 1% low melt agarose and imaged at 40X magnification with a Nikon A1R Si Point Scanning Confocal microscope.

### TurboID affinity purification and mass spectrometry

For TurboID, the affinity purification followed essentially the protocol described in (41). Briefly, embryos were lysed in modified RIPA (modRIPA) buffer [50 mM Tris-HCl, pH 7.4, 150 mM NaCl, 1 mM EGTA, 0.5 mM EDTA, 1 mM MgCl_2_, 1% NP40, 0.1% SDS, 0.4% sodium deoxycholate, 1 mM PMSF and 1x Protease Inhibitor mixture]. The volumes employed were 0.5 µl per embryo for 12 hpf embryos and 2 µL per embryo for 48 and 72 hpf embryos. Embryos were sonicated for 15–30 seconds at 30–35% amplitude (5 seconds on, 3 seconds off for 3 cycles at 30% amplitude for 12 hpf embryos, 10 seconds on, 3 seconds off for 3 cycles at 32% amplitude for 48 hpf embryos, and 10 seconds on, 3 seconds off for 3 cycles at 35% amplitude for 72 hpf embryos) on a Q500 Sonicator with 1/8” Microtip (QSonica, Newtown, Connecticut, Cat# 4422). Following sonication, 250 U of TurboNuclease (BioVision Inc., Milpitas, CA, Cat# 9207) were added and the samples incubated at 4℃ with rotation for 20 minutes. Next 10% SDS was added to the sample to bring the SDS concentration to 0.4%, and the sample incubated at 4 ℃ with rotation for a further 5 minutes. Samples were centrifuged at 16,000 x g for 20 minutes and the supernatant used for affinity purification of biotinylated proteins using 25 µl of washed streptavidin agarose beads (GE Healthcare Life Science, Cat# 17511301) for 16 hours with rotation at 4℃. The beads were next washed once with SDS-Wash buffer (25 mM Tris-HCl, pH 7.4, 2% SDS), twice with RIPA (50 mM Tris-HCl, pH 7.4, 150 mM NaCl, 1 mM EDTA, 1% NP40, 0.1% SDS, 0.4% sodium deoxycholate), once with TNNE buffer (25 mM Tris-HCl, pH 7.4, 150 mM NaCl, 1 mM EDTA, 0.1% NP40), and three times with 50 mM ammonium bicarbonate (ABC buffer), pH 8.0. Proteins were digested on-bead with 1 µg of trypsin (Sigma Aldrich, Cat# 6567) in 50 µL ABC buffer with rotation at 37℃ overnight. A further 0.5 µg of trypsin was added, and digestion continued for 3 hours. Following digestion, the beads were spun down (400 x g) and the supernatant transferred to a new tube. The beads were washed twice with 75 µl water and the wash water collected and added to the supernatant. The pooled supernatant was acidified with 0.1 volume of 50% formic acid and dried by vacuum centrifugation. Peptides were resuspended in 5% formic acid and stored at −80℃.

### Mass spectrometry acquisition

Digested peptides were analyzed by nanoLC-MS/MS in a Data Dependent Acquisition (DDA) workflow. Tryptic peptides from 200 embryos (at 48 and 72 HPF) or 900 embryos (at 12 HPF) were used, and 2/3 of each sample was injected for DDA analysis. Packed column emitters with a 5–8 µm tip opening were generated from 75 µm internal diameter fused silica capillary tubing, using a laser puller (Sutter Instrument Co., model P-2000). Nano-spray emitters were packed with C18 reversed-phase material (Reprosil-Pur 120 C18-AQ, 3 µm) resuspended in methanol using a pressure injection cell to a packed length of 15 cm. Samples were analyzed using an Eksigent 425 nanoHPLC (Redwood, CA) connected to a Thermo Fusion Lumos Mass Spectrometer (San Jose, CA). Samples were loaded directly onto the packed tip emitter using the autosampler at a flow rate of 400 nl/minute. Peptides were eluted from the column using a 90 minutes gradient (2–30% acetonitrile) at 200 nl/minute. After 90 minutes, the organic concentration was increased to 80% over 10 minutes, held for 5 minutes and then the column was equilibrated at 2% acetonitrile for 30 minutes. The DDA method used a 240k resolution and 5e5 target setting for MS1 scans. A 3 seconds cycle was used for HCD MS/MS acquired with 1 Da isolation, 15k resolution, 2e5 target, and 50 ms max fill time with a dynamic exclusion set to 12 seconds.

### Mass spectrometry data analysis

Mass spectrometry data generated were stored, searched and analyzed using ProHits laboratory information management system (LIMS) platform (42). Within ProHits, ProteoWizard (V3.0.1072) (43) was used to convert the .RAW files to .mgf and .mzML formats. The data files were searched using Mascot (V2.3.02) (44) and Comet (V2016.01 rev.2) (45) against zebrafish sequences from the RefSeq database (version 65, July 2nd, 2014), supplemented with “common contaminants” from the Max Planck Institute (http://lotus1.gwdg.de/mpg/mmbc/maxquant_input.nsf/7994124a4298328fc125748d0048fee2/$FILE/contaminants.fasta) and the Global Proteome Machine (GPM; ftp://ftp.thegpm.org/fasta/cRAP/crap.fasta), sequence tags (BirA, GST26, mCherry and GFP), LysC, and streptavidin. Reverse (decoy) entries were created for all sequences for a total of 86,384 entries. Search parameters were set to search for tryptic cleavages, allowing 2 missed cleavage sites per peptide a mass tolerance of 12 ppm and a tolerance of 0.15 amu for fragment ions. Variable modifications included deamidation (asparagine and glutamine) and oxidation (methionine). Results from each search engine were analyzed through TPP (the Trans-Proteomic Pipeline, v.4.7 POLAR VORTEX rev 1) (46) via the iProphet pipeline (47). All proteins with an iProphet probability ≥ 95% and two unique peptides were used for analysis.

### Interaction scoring

SAINTexpress (Version 3.6.1.) (48) was used to score the probability that identified proteins were enriched above background. SAINTexpress uses a model whereby each prey identified in a TurboID experiment with a given bait, is compared using spectral counts as a measure of abundance, against a set of negative controls. For each separate experiment (different bait or condition) independent biological replicates were used to generate the prey profile and this profile was compared against negative controls. A minimum of two biological replicates were used for all analysis. When three purifications were performed for a given bait and conditions, two virtual replicates were generated prior to SAINTexpress scoring. The virtual replicates were generated by selecting the two highest spectral counts for each prey in the purification of the bait across the three replicates. Creating these virtual replicates increases sensitivity when the reproducibility across the biological replicates is not perfect. Negative controls consisted of purifications from embryos expressing TurboID-GFP (6 and 3 replicates for the TurboID mRNA injection and transgenic experiments, respectively) or miniTurbo-GFP (5 and 3 replicates for the miniTurbo mRNA injection and transgenic experiments, respectively) and from embryos not expressing any exogenous fusion proteins (3 replicates for all experiments). For running SAINTexpress, the 6–9 negative controls were compressed to 4 virtual controls, meaning that the four highest spectral counts for each identified protein were used in the scoring to increase stringency. Scores were averaged across both biological replicates (or virtual replicates), and these averages used to calculate a Bayesian False Discovery Rate (BFDR); preys detected with a BFDR of ≤ 1% were considered high-confidence.

### Human orthology

To compare our zebrafish data with human data we converted the zebrafish protein names to their human orthologs. We used the g:profiler g:Convert tool (49) and manual curation from NCBI (50) to make the conversions.

### Gene ontology enrichment analysis

Gene ontology (GO) enrichment analysis was performed using the human orthologs of the identified high-confidence proximal interactors as inputs. The g:Profiler g:GOSt tool (49) was used for the enrichment analysis. Default options were used, with the exceptions that the data source was set to only “GO cellular component terms”, with term size between 5–500.

### Comparison with previously published datasets

To compare our data with previously published datasets, we obtained the lists of high-confidence LMNA proximal interactors from the supplemental data of the Samavarchi-Tehrani et al. (51) and May et al. (52) publications. Using these lists of high-confidence proximal interactors, we performed gene ontology enrichment analysis in the same manner as for our own data (see above). For comparison to the interactors listed on BioGRID, the full curated list of interactors for the *H. sapiens* LMNA protein was obtained from the BioGRID website (https://thebiogrid.org/110186/summary/homo-sapiens/lmna.html, accessed April 04 2021) and compared against the high-confidence proximal interactors identified our experiments.

### Data Visualization

Dot plots, scatter plots, and Pearson correlation data were generated using ProHits-viz (53). In the ProHits-viz dot plots tool, once a prey passes the selected FDR threshold (≤ 1% used here) with one bait, all its quantitative values across all baits are recovered. The FDR of the prey is then indicated by the edge color. Quantitative information (spectral count) is represented by the color gradient within the node, while relative prey counts across all baits are denoted by the size of the node (i.e. the maximal size for each prey corresponds to the bait in which it was detected with the highest number of spectra).

### Experimental Design and Statistical Rationale

For each TurboID experiment, independent biological duplicates or triplicates were used. Each replicate was generated through independent mRNA injections or egg lays (for transgenic experiments) and collections. Statistical scoring using SAINTexpress against 6–9 controls compressed to 4 virtual controls was performed as described above (“Interaction scoring”). The control samples used were TurboID-GFP or miniTurb-GFP matched to the enzyme and expression method used in the experiment, and wild type embryos not expressing exogenous protein, used in all experiments and age / treatment matched to the experiment unless otherwise noted. The same wild type control runs were used in both TurboID and miniTurbo analysis. The Bayesian FDR was calculated using the average SAINTexpress score across replicates. The SAINTexpress analysis was performed independently for each enzyme and expression method. In other words, each of the following sets was analyzed independently: a) the mRNA injection TurboID-LMNA 12 and 48 hour labeling runs, b) the mRNA injection miniTurbo-LMNA 12 and 48 hour labeling runs, c) the transgenic TurboID-LMNA, d) the transgenic miniTurbo-LMNA, and e) the transgenic TurboID-LMNA comparison of labeling conditions.

## Results

### Adapting TurboID and miniTurbo for *in vivo* biotinylation in zebrafish

To determine whether the TurboID and miniTurbo enzymes could efficiently biotinylate proteins in early zebrafish embryos, codon-optimized gene constructs of TurboID and miniTurbo were synthesized for fusion to proteins of interest. The human lamin A (LMNA) bait was selected for optimization experiments as it has a well-characterized proximal interactome in mammalian cells (7, 51, 52) as well as a characteristic subcellular localization, enabling visual evaluation of the functionality of the fusions. This bait was first used to demonstrate the usefulness of the BioID method in identifying the proximal proteome of insoluble proteins that are typically difficult targets for conventional protein-protein interaction methods (7). The human and zebrafish LMNA proteins share a 62% identity and 76% similarity at the amino acid level, and mutations of conserved sites result in similar protein mis-localization in both zebrafish and human cells (54, 55). This suggested that the human LMNA protein would function and localize appropriately when expressed in the zebrafish. As a negative control, the Green Fluorescent Protein (GFP) was subcloned in the same system.

To test the biotinylation efficiency of TurboID and miniTurbo in zebrafish, we used mRNA injections in zebrafish embryos to express TurboID or miniTurbo fused to LMNA or GFP coding sequences. Briefly, capped mRNA for TurboID and miniTurbo fusions were generated by *in vitro* transcription, purified, and injected into single cell stage zebrafish embryos (Figure 1A). The injected embryos were grown in egg water supplemented with biotin for 12 or 48 hours. Labelling was stopped by washing the embryos with ice-cold egg water without biotin. The embryos were dechorionated, deyolked, washed, and frozen for later analysis.

**Figure 1.**
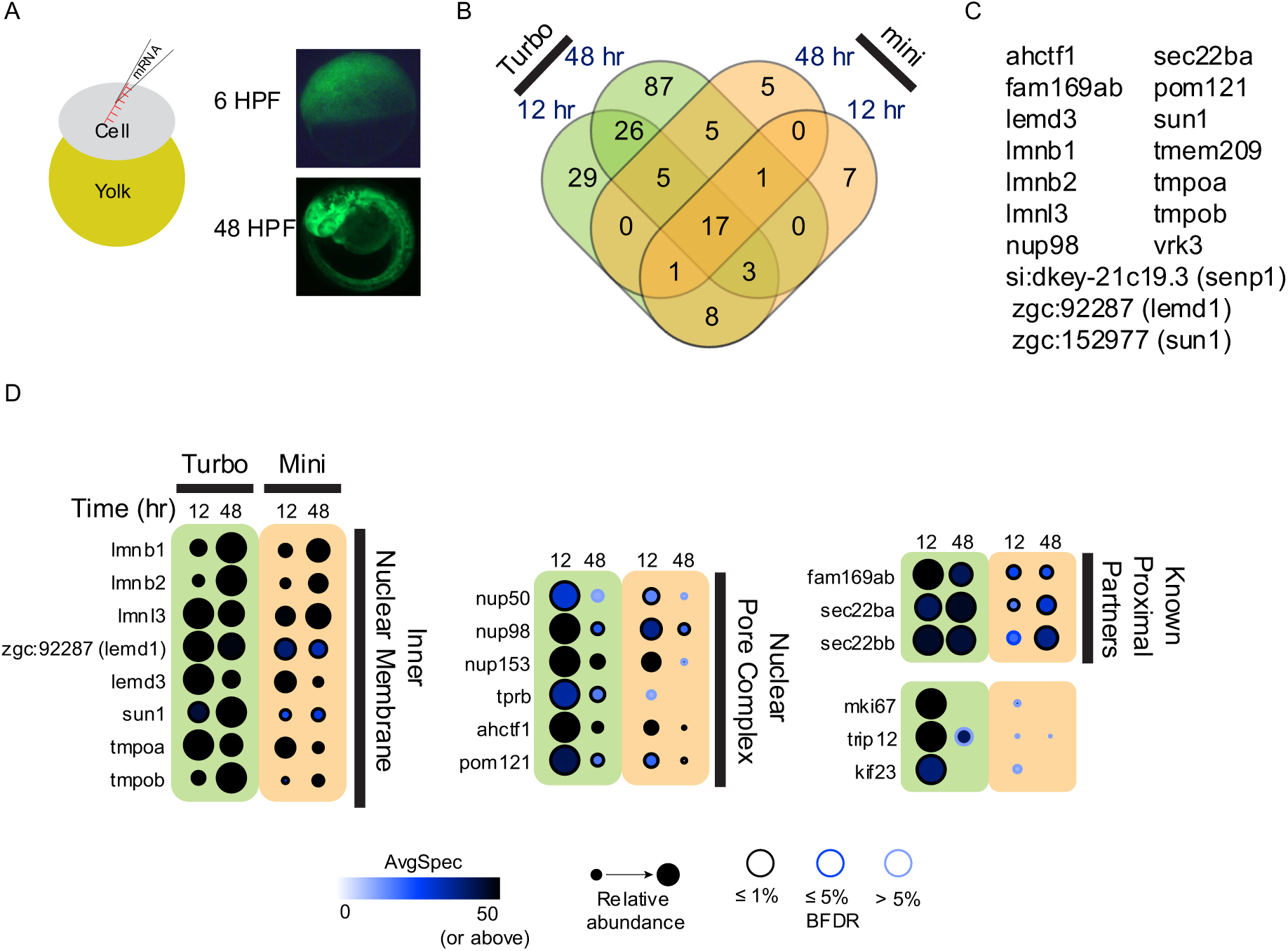
mRNA injection-based expression identifies LMNA proximal proteins in the early zebrafish embryo. (A) *In vitro* transcribed mRNA coding for TurboID fused to GFP was injected into the single cell stage zebrafish embryo and GFP fluorescence was visualized at 6 and 48 hpf (magnification 16x). (B) TurboID-LMNA or miniTurbo-LMNA were expressed by mRNA injections (with the corresponding GFP fusions used as negative controls). Injected embryos were collected at 12 and 48 hpf. The number of proteins detected at BFDR ≤ 1% under each of the conditions is shown. (C) List of the 18 high-confidence proximal interactors shared by all four conditions in (B). (D) The 25 most abundant proteins (by spectral counts) with a BFDR ≤ 1% from the 12 hour TurboID-LMNA condition are listed, alongside their abundance across each of the conditions.

To determine what level of biotin supplementation was required to achieve optimal protein biotinylation, we injected TurboID-LMNA and miniTurbo-LMNA mRNA into 100 embryos per condition and incubated them in egg water supplemented with increasing biotin concentrations (0, 200 and 800 µM) for 48 hours. 50 µg total lysate (equivalent to approximately 10 embryos) per condition were separated onto SDS-PAGE gels and used to assess the extent of total biotinylation using streptavidin-horseradish peroxidase (Strep-HRP) blots. The levels of expressed tagged proteins were detected using an anti-BirA antibody capable of detecting TurboID and miniTurbo fusions. In the absence of any biotinylation enzyme injection (uninjected embryos), several biotinylated bands were readily detected in the absence of biotin, and biotin supplementation increased this signal ∼1.9-fold at the highest concentration used. Injection of TurboID-LMNA (and to a lesser extent, miniTurbo-LMNA) increased the signal up to ∼2.2-fold in the absence of biotin and up to ∼2.7-fold after biotin addition. Many of the bands were in common to all conditions, indicating that they are background proteins (such background proteins have been well-documented in cell culture and notably include endogenously biotinylated proteins such as mitochondrial carboxylases (56, 57)). However, some of the bands appear specific to the TurboID and miniTurbo conditions (Supplemental Figure S1). We also note that the high signal detected with TurboID in the absence of biotin is consistent with reports that it has a high avidity for biotin, and is capable of scavenging even small concentrations of this metabolite (9). We did not notice any obvious developmental delay or morphological deformity with either concentration of biotin or injection of (mini)Turbo fusions, suggesting low toxicity in our system (not shown). We therefore selected the highest (800 µM, near the upper limit of biotin solubility in water) concentration of biotin tested for the next set of experiments.

### Characterization of the lamin A proximal interactome following mRNA injection in zebrafish

To explore whether the signal-to-noise was sufficient to identify proximity partners for LMNA in zebrafish embryos (or whether the high background was limiting the usefulness of the system), we next purified biotinylated proteins via streptavidin pulldown and identified them by mass spectrometry. To assess whether biotinylation was specific for our protein of interest, we compared the proximal proteome of LMNA (fused to either TurboID or miniTurbo) to that of GFP (fused with the same enzymes) and also purified biotinylated proteins from uninjected embryos (WT). Briefly, embryos (900 per replicate for the 12 hour time point, and 200 for 48 hours) were injected with the indicated mRNA. Following injection, embryos were incubated in biotin-supplemented egg water for labeling, for 12 or 48 hours. The 12 hour labeling time covers development through gastrulation to the start of somitogenesis (58), while the 48 hour labeling time covers early development until the hatching stage at the completion of much of primary organogenesis (58). Labeling was stopped by washing embryos in ice-cold egg water. To enrich for biotinylated proteins, streptavidin pulldowns were performed on lysates from pools of mRNA-injected embryos, and non-biotinylated peptides were eluted by tryptic digestion. Released peptides were identified by mass spectrometry, and peptides assigned to proteins by the Trans-Proteomic Pipeline iProphet (iProphet) tool (46, 47). Proteins identified with an iProphet probability ≥ 95% and two unique peptides were used for analysis. Experiments were performed using biological replicates (duplicates for 12 hours and triplicates for 48 hours), with the following exceptions: for the negative control TurboID-GFP four replicates were used, for uninjected embryos only one replicate was used at 12 hours and two at 48 hours. Between 412 proteins (uninjected, 48 hours) and 1279 proteins (TurboID-LMNA, 48 hours) were identified per replicate (Table 1). Across all experiments, the 48 hours labeling yielded a higher number of identified proteins than the 12 hours labeling, as expected. TurboID yielded 23–29% more identifications than the corresponding miniTurbo constructs, with the exception of the 48 hours GFP controls where identifications were almost equal (Table 1). For all labelling times, the proteins detected with the highest spectral counts were shared between the LMNA and GFP baits, suggesting that they are non-specific (background) proteins. At the 12 hour time point, these consisted mainly of mitochondrial and yolk proteins, while at 48 hours yolk protein detection was reduced and muscle proteins became highly abundant.

**Table 1.** Unique proteins identified by mRNA injection based BioID experiments. The number of unique proteins identified by ≥ 2 unique peptides and an iProphet probability ≥ 95% are listed for all mRNA injection-based experiments. For LMNA bait experiments the number of proteins with a SAINT BFDR ≤ 1% are listed.

| Parameter<br>Experiment / Replicate | | Unique<br>Proteins<br>Identified | Total Unique<br>Proteins<br>Identified | Shared<br>Unique<br>Proteins<br>Identified | Proteins with<br>a<br>SAINT BFDR<br>$\leq 1\%$ |
| --- | --- | --- | --- | --- | --- |
| TurboID-LMNA<br>12 Hour Labeling | Rep 1 | 774 | 982 | 616 | 89 |
|  | Rep 2 | 824 |  |  |  |
| TurboID-LMNA<br>48 Hour Labeling | Rep 1 | 1010 | 1708 | 668 | 144 |
|  | Rep 2 | 1279 |  |  |  |
|  | Rep 3 | 1173 |  |  |  |
| miniTurbo-LMNA<br>12 Hour Labeling | Rep 1 | 759 | 854 | 557 | 37 |
|  | Rep 2 | 642 |  |  |  |
| miniTurbo-LMNA<br>48 Hour Labeling | Rep 1 | 970 | 1273 | 495 | 34 |
|  | Rep 2 | 727 |  |  |  |
|  | Rep 3 | 851 |  |  |  |
| TurboID-GFP<br>12 Hour Labeling | Rep 1 | 864 | 1163 | 701 | N/A |
|  | Rep 2 | 1000 |  |  |  |
| TurboID-GFP<br>48 Hour Labeling | Rep 1 | 1278 | 1586 | 492 | N/A |
|  | Rep 2 | 714 |  |  |  |
|  | Rep 3 | 867 |  |  |  |
|  | Rep 4 | 1032 |  |  |  |
| miniTurbo-GFP<br>12 Hour Labeling | Rep 1 | 751 | 927 | 628 | N/A |
|  | Rep 2 | 804 |  |  |  |
| miniTurbo-GFP<br>48 Hour Labeling | Rep 1 | 1076 | 1525 | 565 | N/A |
|  | Rep 2 | 756 |  |  |  |
|  | Rep 3 | 1199 |  |  |  |
| Uninjected<br>12 Hour Labeling | Rep 1 | 621 | N/A | N/A | N/A |
| Uninjected<br>48 Hour Labeling | Rep 1 | 412 | 844 | 325 | N/A |
|  | Rep 2 | 757 |  |  |  |

We next applied SAINTexpress (SAINT) (48) to identify proteins specific to the LMNA fusion conditions. SAINTexpress analysis of this dataset was performed as follows. First, for the 48 hour labeling, we combined the triplicate purifications of the baits into two virtual replicates (i.e. the two highest spectral counts for each prey in the purification of the bait across the three replicates are selected); this increases sensitivity when the reproducibility across the biological replicates is not perfect (see Experimental Procedures). We also combined using the same approach the eight (for miniTurbo) or nine (for TurboID) negative controls into to four virtual controls (as in (59)); this step increases the stringency in SAINTexpress scoring. The controls consisted of uninjected embryos (duplicate of 48 hpf embryos and one run of 12 hpf embryos) and six or five replicates of TurboID or miniTurbo, respectively (two replicates at 12 hpf for both enzymes, four and three replicates at 48 hpf for TurboID-GFP and miniTurbo-GFP respectively). All negative control samples were incubated with 800 µM biotin for the duration of the experiment. We are considering as high-confidence proximal partners those that pass a Bayesian False Discovery Rate (BFDR) cutoff of ≤ 1%. This revealed 89 and 144 high-confidence proximal interactors identified at 12 and 48 hours, respectively, for TurboID-LMNA and 37 and 34 proximal interactors with miniTurbo-LMNA (Table 1; Supplementary Table 1). Of these unique proteins, 40 were identified by both TurboID-LMNA and miniTurbo-LMNA (at either time points; Figure 1B), with 17 identified across all conditions (Figure 1C). The 20 proteins with the highest spectral counts and a BFDR ≤ 1% in the TurboID-LMNA 12 hour labeling condition included proteins known to localize to the inner nuclear membrane, components of the nuclear pore complex, and previously identified LMNA proximal partners (Figure 1D). From this list, all the proteins annotated to the inner nuclear membrane were readily detected across all conditions. The nuclear pore complex proteins displayed greater differences across conditions, with the 48 hour labeling time producing a lower relative abundance with either TurboID or miniTurbo (Figure 1D).

Correlation between replicates is an important metric of the reproducibility of the labeling experiments and is an important component of the SAINT scoring (i.e. only preys enriched across replicates are considered significant). For post SAINT correlations, we used a BFDR ≤ 5% as a cutoff, to include proteins just missing the ≤ 1% BFDR cutoff. The 12 hour labeling time resulted in very high correlation between replicates, with pairwise correlation pre and post SAINT ranging from 0.87–0.98 (Figure S2). As expected (due to the increased complexity of the tissue and the longer labeling time allowing labeling of more background proteins), with 48 hours of labeling there was greater variability, with pairwise correlation between replicates lower than with 12 hours of labeling (range 0.36–0.89; Figure S3). We conclude that for most applications, 12 hours of labeling is sufficient and will allow for easier filtering, using fewer control samples. 48 hour labeling produces higher background, but not significantly higher labeling of proximal interactors, thereby reducing the overall sensitivity after background contaminant filtering.

Taken together, these results indicate that, using mRNA injection as an expression method, TurboID-LMNA and miniTurbo-LMNA specifically biotinylate proximal proteins at a level high enough to allow for identification above background.

### Generation of inducible transgenic zebrafish lines for *in vivo* TurboID

Having shown that proximity dependent biotinylation and proximal protein identification could be achieved in zebrafish embryos, we progressed to performing these experiments in inducible stable transgenic zebrafish. This method has the advantage of allowing control of labeling via inducible expression of the transgene at any timepoint during development. To allow for temporal control of expression, we selected the *hsp70l* heat shock inducible promoter, which is well tolerated by zebrafish embryos (60), and does not require drug treatment. Using the Tol2 transgenesis system, stable transgenic zebrafish were generated to express, under the control of the 1.5kb *hsp70l* promoter, TurboID or miniTurbo fused to either LMNA or GFP as a control, resulting in the generation of four lines. All protein fusions included a 3X FLAG tag in addition to the biotin ligase to enable detection of the expressed protein and immunoprecipitation. We designate these transgenic lines: *Tg(hsp70l:TurboID-LMNA)^hsc158^*, *Tg(hsp70l:TurboID-GFP)^hsc159^*, *Tg(hsp70l:miniTurbo-LMNA)^hsc160^*, and *Tg(hsp70l:miniTurbo-GFP)^hsc161^*. All transgenic constructs also included GFP expressed under the control of the cardiomyocyte specific *myl7* promoter (34), allowing for the identification and selection of integration-positive embryos by visual inspection of robust GFP fluorescence in the heart, generally assessed at 48 hpf, but visible as early as 16 hpf (Figure 2A).

**Figure 2.**
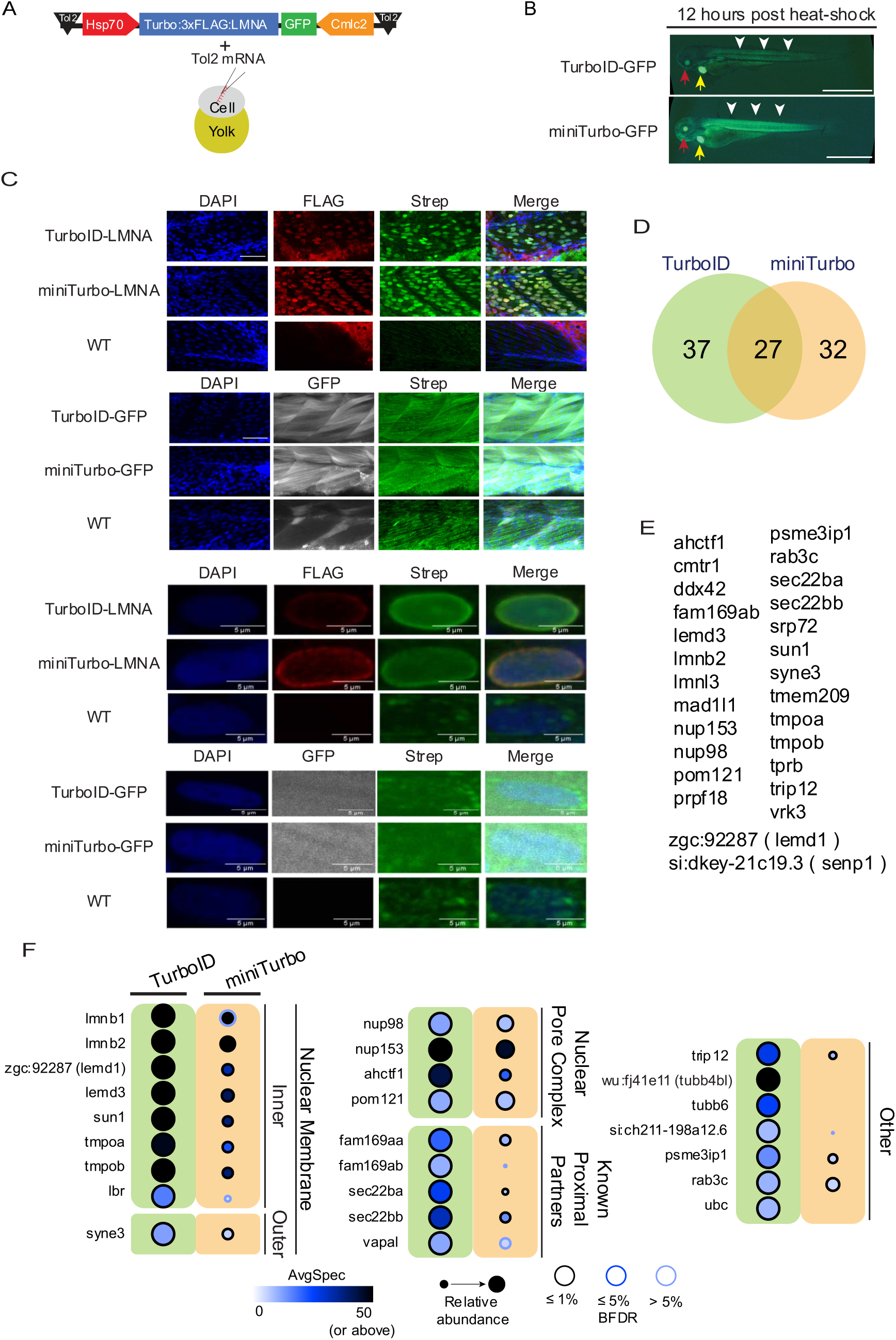
Inducible transgenic expression of TurboID-LMNA or miniTurbo-LMNA identifies LMNA proximal proteins. (A) Cassettes compatible with the Tol2 trangenesis system were generated Proximity-dependent biotinylation in zebrafish embryos that express the selected fusion proteins (TurboID-LMNA and GFP, miniTurbo-LMNA and GFP) downstream of the heat shock inducible Hsp70 promoter. The cassettes also contain GFP under the control of a cardiomyocyte specific Cmlc2 promoter to allow selection of integration-positive embryos. (B) TurboID-GFP and miniTurbo-GFP transgenic embryos were heat shocked at 60 hpf for one hour and imaged at 72 hpf. Integration-positive embryos were selected prior to heat shock based on expression of GFP in the heart (yellow arrow). GFP expression 12 hours following heat shock is indicated by white arrowheads, and GFP expression in the eye lens by red arrows (magnification 16x, Scale Bar = 1 mm). (C) Immunofluorescence microscopy on embryos collected at 72 hpf following 1 hr heat shock at 60 hpf time point was performed as detailed in Experimental Procedures FLAG staining to detect the bait expression and streptavidin-fluorophore (Strep) to detect biotinylation. DAPI was used to stain the nucleus. (Upper panels Scale bar 50 µm, magnification 20x; Lower panels scale bar 5 µm, magnification 40x). (D) Number of high-confidence proximity partners for TurboID and miniTurbo, respectively. (E) List of the 27 common partners. (F) The 25 most abundant proteins (by spectral counts) with a BFDR ≤ 1% from the TurboID-LMNA experiment are listed, alongside their abundance across both conditions.

Expression was induced by heating zebrafish embryos to 38℃ for one hour prior to returning the embryos to their normal incubation temperature (28.5℃) for 11 hours. The heat shock induced low levels of visible fluorescence by three hours post-initiation, as visualized by TurboID-GFP fluorescence, with robust ubiquitous expression visible from five hours through to collection of embryos at twelve hours post heat shock. GFP expression is expected in the eye even without heat shock since the *hsp70l* promoter endogenously drives expression in the eye lens during early development (61) (Figure 2B).

To ensure that the fusion proteins were localizing correctly and that protein biotinylation was occurring specifically proximal to the fusion protein, whole mount immunofluorescence was performed to determine the localization of TurboID-LMNA, miniTurbo-LMNA, TurboID-GFP, miniTurbo-GFP and to define where biotinylation occurred. An anti-FLAG antibody was used to visualize the fusion proteins and streptavidin conjugated to a fluorophore was employed to visualize biotinylation. Embryos were incubated in water supplemented with 800 µM biotin, or no biotin as a control. Protein expression was induced by heat shock at 60 hpf for 1 hour at 38℃. Following heat shock, the embryos were incubated for 12 hours before collection at 72 hpf. Immunofluorescence showed that TurboID-LMNA and miniTurbo-LMNA are localized, as expected, around the nucleus. Biotinylated proteins were also localized surrounding the nucleus, suggesting that biotinylation was occurring specifically proximal to the fusion proteins (note that this biotinylation signal may include the bait itself). This contrasted with TurboID-GFP and miniTurbo-GFP where GFP and biotinylation were visible throughout the cell. In all samples, background streptavidin staining was evident in the mitochondria (Figure 2C).

### Inducible expression of TurboID and miniTurbo in transgenic zebrafish embryos can be used to identify proximal partners

Having shown that the inducible fusion protein is properly localize (and leads to biotinylation in the expected locale), we moved to performing TurboID-based proteomics in transgenic zebrafish embryos. We wanted to utilize the inducible transgenic lines to perform TurboID experiments outside the time window possible with mRNA injections. For this reason, we elected to induce transgene expression at 60 hpf. We also selected the shortest labeling time we had tested with mRNA injection, 12 hours, as this short labeling time would demonstrate the utility of this system for tracking developmental processes over time. This labeling period corresponds to the final stages of zebrafish embryonic growth, with the larval stage beginning at 72 hpf (58).

For all experiments, protein expression was induced by heat shock of the embryos at 38℃ for one hour. Embryos expressing TurboID and miniTurbo fused to GFP and wild type embryos were used as controls. The SAINTexpress software (SAINT) was used to filter the identified protein lists. By comparing against the control protein lists and between replicates, filtered lists of proteins unique to the experimental condition were generated.

For our initial transgenic TurboID experiments, TurboID-LMNA and miniTurbo-LMNA expression was induced at 60 hpf, labeling continued for 12 hours, and the embryos collected and labeling terminated at 72 hpf. The embryos were incubated in egg water supplemented with 800 µM biotin for the full 72 hours. This prolonged biotin supplementation prior to induction of transgene expression was selected to ensure sufficient time for biotin accumulation within the embryo, as some polar compounds can require extended incubation to achieve high internal concentrations in zebrafish embryos (62). Following affinity purification and mass spectrometry, a total of 662 and 692 proteins were identified across triplicates for TurboID-LMNA and miniTurbo-LMNA respectively (Table 2).

**Table 2.**
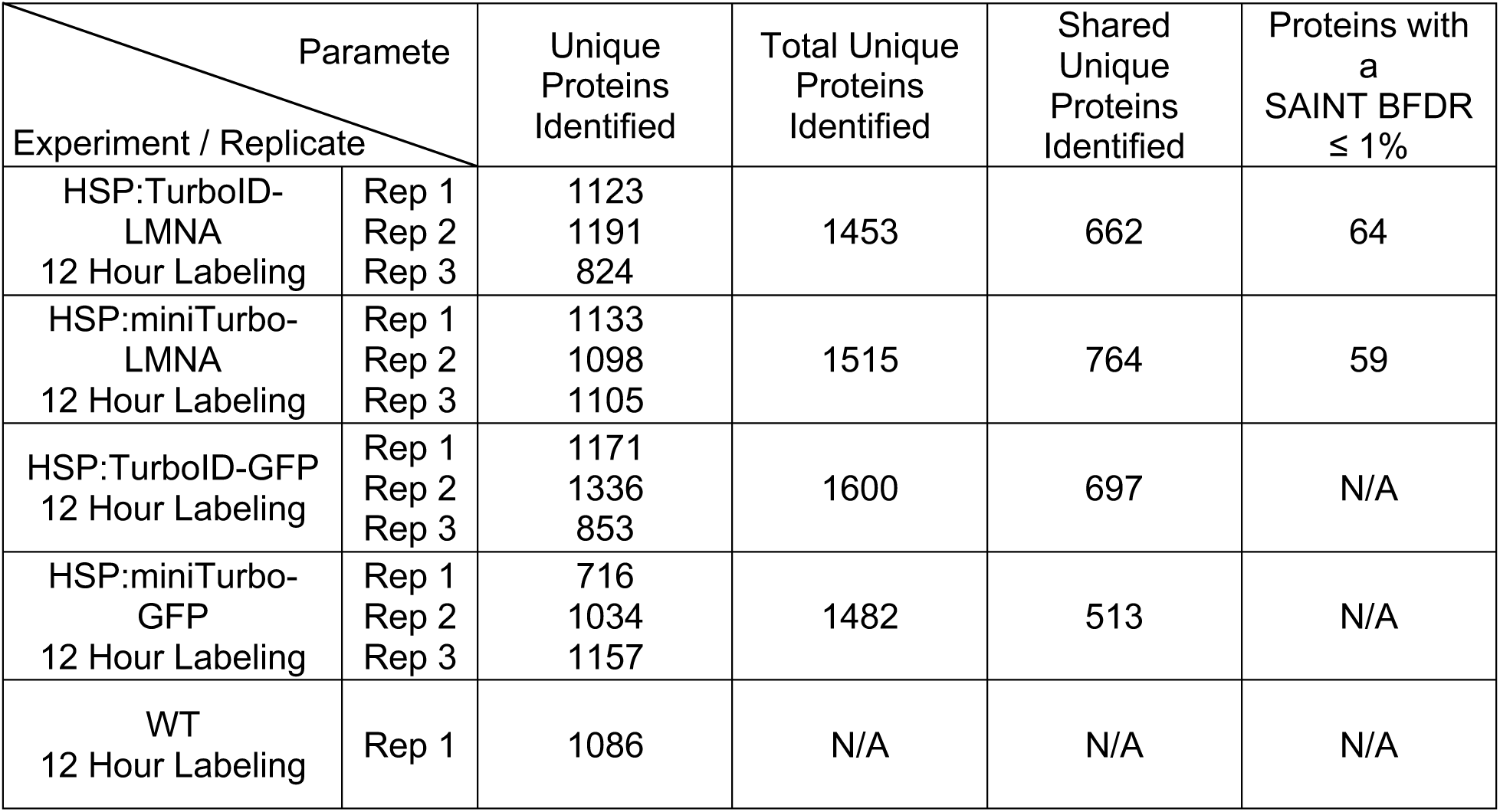
Unique proteins identified by heat shock inducible transgenic based BioID experiments. The number of unique proteins identified by ≥ 2 unique peptides and an iProphet probability ≥ 95% are listed for all heat shock inducible transgenic experiments. All experiments used one hour heat shock (38℃) induction at 60 hpf, and collection at 72 hpf. For LMNA bait experiments the number of proteins with a SAINT BFDR ≤ 1% are listed.

SAINTexpress analysis of this dataset was performed as for the mRNA injection dataset, with the triplicates combined into two virtual replicates and the six controls combined into four virtual controls. The controls consisted of wild type embryos (duplicate of 48 hpf embryos and one run of embryos heat shocked at 60 hpf and collected at 72 hpf) and triplicates of TurboID or miniTurbo fused to GFP (heat shocked at 60 hpf and collected at 72 hpf). All negative control samples were incubated with 800 µM biotin for the duration of the experiment. Correlation between replicates ranged from 0.27–0.80 prior to SAINTexpress scoring and 0.36–0.81 when considering only hits with a BFDR ≤ 5% (Figure S4). Following SAINTexpress scoring, 64 and 59 proteins with a BFDR ≤ 1% remained for TurboID-LMNA and miniTurbo-LMNA, respectively, with 27 of them shared (Figure 2D and E; Supplementary Table 2). Seventeen of these 27 shared proteins were among the 25 most abundant (after SAINT filtering) high-confidence proximal interactors (by spectral count) identified by TurboID-LMNA with a BFDR ≤ 1% (Figure 2F).

### Gene Ontology enrichment analysis

To determine if the identified high-confidence proximal interactors were enriched for proteins known to localize near LMNA, we performed Gene Ontology (GO) enrichment analysis. We first converted the zebrafish genes to their human orthologs to take advantage of the greater degree of annotations (Supplementary Table 3). GO profiling of the human orthologs of the identified proximal interactors with a BFDR ≤ 1% found “Nuclear Membrane” and “Nuclear Envelope” to be the most highly enriched Cellular Component (CC) categories (CC categories containing 5–500 terms). This was true for both TurboID and miniTurbo in both mRNA injection and transgenic experiments (Supplementary Table 4). For mRNA injection-based experiments, between 18 and 32 of the high-confidence proximal interactors were annotated with these terms. For TurboID-LMNA and miniTurbo-LMNA inducible transgenic TurboID experiments, these terms annotated 26 and 18 high-confidence proximal interactors, respectively (Table 3, these numbers are higher than the direct output from g:Profiler since some annotated genes are duplicated in the zebrafish genome and both proteins were detected), with 14 of them detected with both enzymes. These results demonstrate that both mRNA injection-based and inducible transgenic expression of TurboID-LMNA and miniTurbo-LMNA can specifically biotinylate proximal proteins at a sufficiently high level as to achieve consistent identification above background (Figure 3A and B). A large proportion (24–55%) of the high-confidence proximal interactors in all experimental conditions are annotated as nuclear envelope or nuclear membrane proteins, supporting the conclusion that the labeling is occurring specifically proximal to the bait (Table 3).

**Table 3.**
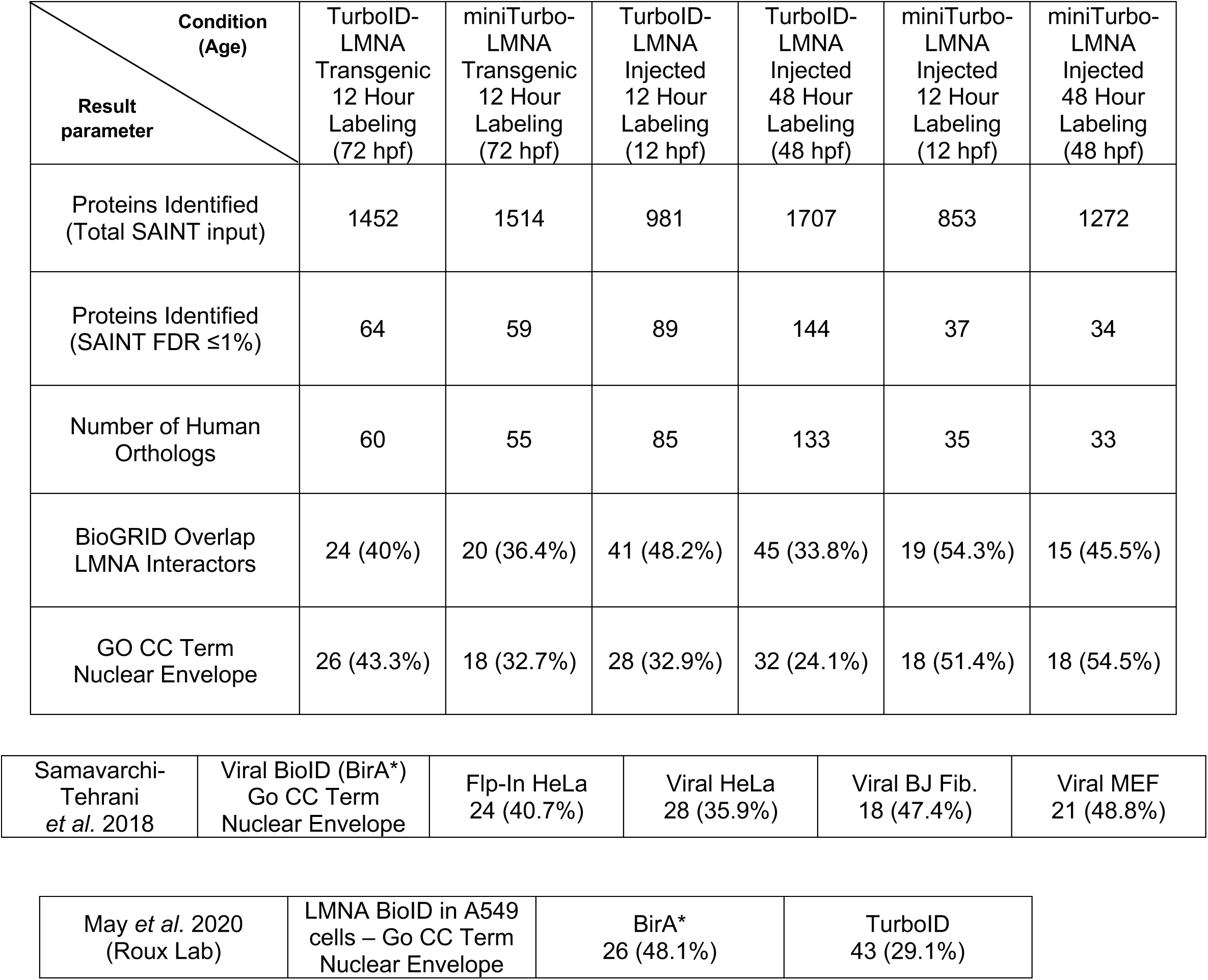
Summary of protein identifications for all zebrafish BioID experiments and comparison to previously published datasets. The total number of proteins identified in each BioID experiment is listed together with the number of high confidence interactors, human orthologs, and number annotated with the GO CC terms “Nuclear Envelope”. The number of genes with this annotation from the Samavarchi-Tehrani *et al.* and May *et al.* publications are also listed.

| Condition (Age)<br>Result parameter | TurboID-LMNA Transgenic 12 Hour Labeling (72 hpf) | miniTurbo-LMNA Transgenic 12 Hour Labeling (72 hpf) | TurboID-LMNA Injected 12 Hour Labeling (12 hpf) | TurboID-LMNA Injected 48 Hour Labeling (48 hpf) | miniTurbo-LMNA Injected 12 Hour Labeling (12 hpf) | miniTurbo-LMNA Injected 48 Hour Labeling (48 hpf) |
| --- | --- | --- | --- | --- | --- | --- |
| Proteins Identified (Total SAINT input) | 1452 | 1514 | 981 | 1707 | 853 | 1272 |
| Proteins Identified (SAINT FDR $\leq 1\%$) | 64 | 59 | 89 | 144 | 37 | 34 |
| Number of Human Orthologs | 60 | 55 | 85 | 133 | 35 | 33 |
| BioGRID Overlap LMNA Interactors | 24 (40%) | 20 (36.4%) | 41 (48.2%) | 45 (33.8%) | 19 (54.3%) | 15 (45.5%) |
| GO CC Term Nuclear Envelope | 26 (43.3%) | 18 (32.7%) | 28 (32.9%) | 32 (24.1%) | 18 (51.4%) | 18 (54.5%) |

|  |  |  |  |  |  |
| --- | --- | --- | --- | --- | --- |
| Samavarchi-Tehrani *et al.* 2018 | Viral BioID (BirA*)<br>Go CC Term<br>Nuclear Envelope | Flp-In HeLa<br>24 (40.7%) | Viral HeLa<br>28 (35.9%) | Viral BJ Fib.<br>18 (47.4%) | Viral MEF<br>21 (48.8%) |

**Table 3. Summary of protein identifications for all zebrafish BioID experiments and comparison to previously published datasets.**
|  |  |  |  |
| --- | --- | --- | --- |
| May *et al.* 2020<br>(Roux Lab) | LMNA BioID in A549 cells – Go CC Term<br>Nuclear Envelope | BirA*<br>26 (48.1%) | TurboID<br>43 (29.1%) |

**Figure 3.**
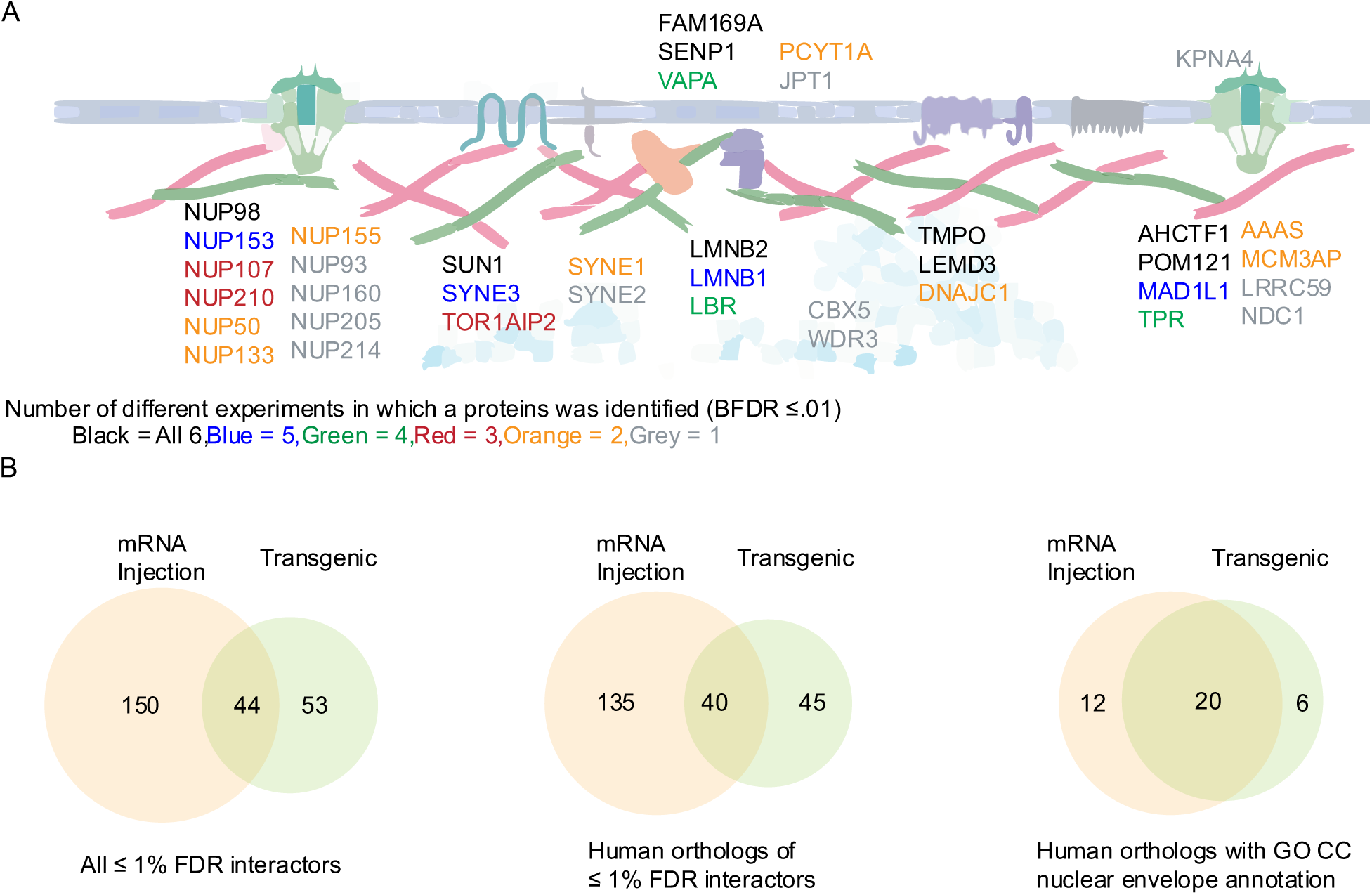
Functional enrichment of LMNA proximal partners. (A) Schematic showing all the high-confidence interactors annotated with the term nuclear envelope or nuclear membrane. The color coding indicates in how many of the experiments this interactor was identified. (B) Overlap between all high-confidence proteins identified using both TurboID and miniTurbo by mRNA injection or transgenic expression (left), their mapped human orthologs (middle), and the human orthologs annotated as “nuclear envelope” by GO CC (right).

### Comparison with other datasets

To further benchmark our data, we compared the human orthologs of the identified zebrafish proteins with published lamin A interactor data. The BioGRID database lists 893 (*H. sapiens*, accessed April 04 2021) unique interactors for the human lamin A (LMNA) protein. These interactors were identified predominantly by yeast two-hybrid (∼ 38%), affinity capture-MS (∼ 25%) and proximity labeling (∼ 25%). Across the different conditions we tested, 41–58 % of the proximal interactors identified were already annotated in the BioGRID database (Table 3).

We next compared our data to published BioID data sets which used LMNA as a bait. We selected these because they allow us to compare specifically to datasets from our own laboratory (i.e. generated with consistent purification and spectrometric analysis protocols (51)) or using TurboID (52). Samavarchi-Tehrani *et al*. (51) used LMNA as a bait in their development of a lentiviral BioID toolkit and profiled LMNA in four conditions. In their experiments, 18–28 (36–49 % of the total high-confidence interactors identified) of the proteins identified in various cell lines with a BFDR ≤ 1% had GO CC annotations of nuclear membrane or nuclear envelope. More recently, May *et al.* (52) identified 26 and 43 proteins (48 and 29 % of the total high-confidence interactors identified) with these annotations in BioID experiments using BirA* and TurboID, respectively, in cell culture. Our results with 18–32 (25–49 % of the total high-confidence interactors identified) filtered hits having these annotations demonstrates that our method can produce results on par with previous data (Table 3). We then compared all the high-confidence hits across these three datasets (all conditions were pooled for each dataset). Our zebrafish dataset shared 32 and 31 hits with the Samavarchi-Tehrani *et al*. and May *et al.* datasets, respectively, while these datasets shared 40 hits with each other, 22 of which were among those also in common to our zebrafish dataset (Supplementary Table 5). We conclude that our zebrafish dataset identifies similar numbers and percentages of known nuclear envelope proteins as identified in previously published datasets, demonstrating that performing proximity labeling in zebrafish using our TurboID-based method is a robust method for identifying proximal interactors *in vivo* in a vertebrate animal model.

### Parameters influencing the recovery of proximal partners in transgenic experiments

We next tested, using the TurboID transgenics system, the effect of reducing the concentration of biotin, omitting the heat shock, and omitting the pre-incubation step with biotin. Omitting biotin or using concentrations < 800 µM drastically reduced labeling efficiency. Summed spectra (across 2 replicates) for the 54 proteins identified by TurboID-LMNA with 800 µM biotin added at 0 hpf with a BFDR ≤ 1% ranged from 2351 counts with 800 µM added at 0 hpf to 337 with no biotin (Figure 4A; Supplementary Table 6). Without induction of protein expression by heat shock, the total spectral counts for these 54 TurboID-LMNA high-confidence proximal interactors were further reduced (Figure 4A; Supplementary Table 6), and no protein passed the SAINT filtering threshold (Figure 4B; Supplementary Table 7). The slightly higher level of spectral counts compared to control (Figure 4A) may be due to the expression of HSP70 in the zebrafish lens during normal development (61) that drives transgene expression for a short period even in the absence of heat shock induction; alternatively, low level leaky expression of the transgene could explain these results (63, 64). Regardless, the low level of capture confirmed that the system is sufficiently non-leaky and that the induction timing can be controlled.

**Figure 4.**
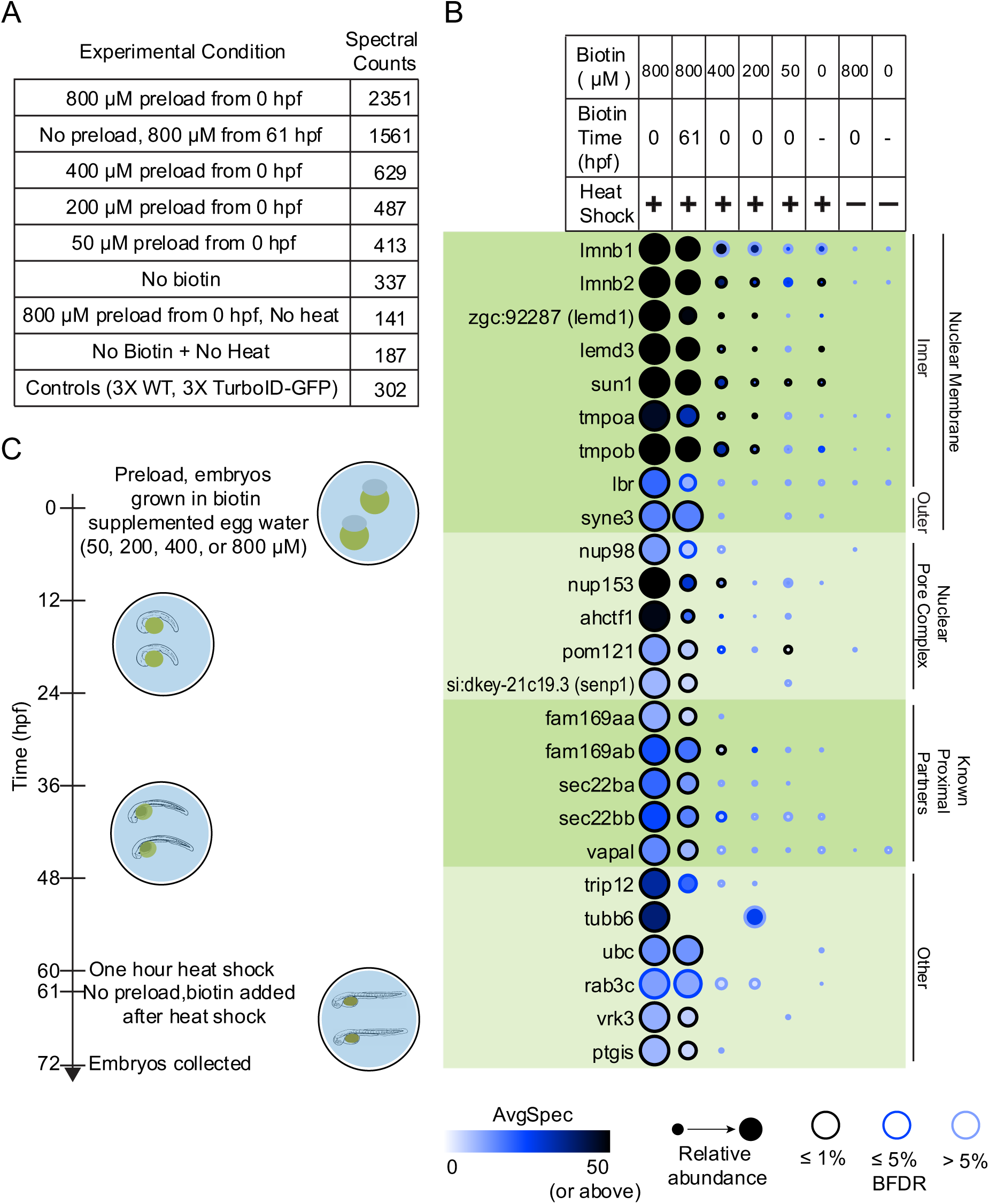
Parameters influencing the recovery of proximal partners in transgenic experiments. (**A**) The total spectral counts from both replicates for the 54 proteins identified with a BFDR ≤ 1% in the 800 µM preload condition were summed for each condition and the total compared. (**B**) The 25 most abundant proteins (by spectral counts) with a BFDR ≤ 1% from the earlier TurboID-LMNA Proximity-dependent biotinylation in zebrafish embryos experiment (Figure 2F) are listed, alongside their abundance across conditions. (C) Schematic of the timeline for biotin supplementation, induction of expression, and collection of embryos.

To see whether the preloading with biotin that we had performed was necessary to enable efficient labeling, we performed an experiment in which biotin was added to the egg water only following heat-shock, from 61–72 hpf (rather than being present from 0–72 hpf) (Figure 4C). In this case, 953 proteins were identified across replicates, compared to 941 for the 0–72 hpf supplementation (this number is lower than earlier as two replicates were used for this comparison vs. three earlier). After filtering against controls using SAINT, 56 and 54 proteins with an BFDR ≤ 1% remained for the 61 hpf and 0 hpf biotin supplementation samples, respectively, of which 32 overlapped (Supplementary Figure 5 A-C; Supplementary Table 7). The shared proteins between these conditions were largely among the more abundant (by spectral count) high-confidence interactors, with 21 of the shared interactors being among the top 25 most abundant interactors from each condition. GO profiling of the human orthologs of the identified high-confidence interactors found 23 and 22 proteins annotated as nuclear envelope for the 0–72 and 61–72 hour biotin supplementation conditions respectively, 19 of which were shared (Supplementary Table 8). These results indicate that omitting the preincubation with biotin still enables recovery of relevant proximal partners, albeit at a slightly lower level.

## Discussion

In this study, we have optimized two methods, mRNA injection and transgenesis, for performing proximity dependent TurboID labeling in zebrafish embryos.

Two recent publications have already demonstrated the use of proximity-dependent labeling in zebrafish. Xiong *et al*. (32) used a conditionally stabilized GFP-nanobody fused to TurboID to target proximal labeling to GFP tagged proteins in muscle, while Pronobis *et al*. (26) used BioID2 targeted cardiomyocytes to investigate proteome changes during cardiac regeneration in adult fish. Both these studies showed promise for investigating proximal interactors *in vivo* in the zebrafish, and the Xiong *et al*., manuscript is particularly interesting in that it permits use of existing green fluorescent protein fusion transgenics directly after a cross with a universal GFP-nanobody expressing line (32). Our manuscript differs from these two studies in a number of ways. Firstly, we determined that both miniTurbo and TurboID are appropriate for proximity-dependent biotinylation in zebrafish. Secondly, we optimized both an mRNA injection and a transgenic system for proximity-dependent biotinylation, increasing the flexibility of the zebrafish biotinylation toolbox (discussed below). Thirdly, we performed an extensive comparison with previously published proximity labeling datasets in cell lines, making the assessment of the robustness of the methods more comprehensive. Lastly, using TurboID, we were able to perform proximity labeling in shorter labeling times (12 hours) than what was published previously, which should enable better temporal studies of developmental stages.

The greatest advantage of using mRNA injection-based expression is that it allows interrogating the proximal proteome of a protein of interest in days rather than in the weeks/months required for the generation of transgenic animals. Protein expression in mRNA injected embryos is ubiquitous and robust, and we show it can be used to reliably identify proximal interactors. While the need to inject hundreds of embryos per replicate can seem daunting, we have not found this to be an obstacle, as we regularly inject 1000+ embryos in a single morning. Additionally, embryos can be frozen following deyolking and pooled from multiple injections to generate sufficient quantity for analysis. We also note that further advances in methods to purify biotinylated proteins from small amounts of input material (65) and in sensitive mass spectrometers (e.g. (66)), may further decrease the workload associated with mRNA injection. Another advantage is the ease with which protein expression levels can be varied, simply by altering the mRNA concentration injected. Some drawbacks of mRNA injection-based expression are the relatively short duration of expression, inability to control expression timing, and inability to restrict expression to specific tissues or cell types. Ultimately, whether mRNA injection is the method of choice depends on the question asked. If the experimental goal is simply to identify the proximity interactome of a protein of interest during early development, and the specific timing of the labeling is not critical, mRNA injections will be a fast and effective choice. Additionally, mRNA injections can be performed in embryos from transgenic crosses, allowing for the comparison of proximity interactomes across genetic backgrounds, such as knockouts, without the lengthy process of generating multiple new transgenic crosses for each background. In some instances, researchers may also prefer to initially perform their proximity labeling using mRNA injections, before moving on to generating stable transgenic lines. This would at the minimum provide some evidence that the tagging of the protein of interest with the enzyme does not alter its localization and permits recovery of specific biotinylated preys.

On the other hand, while generating transgenic animals requires a greater upfront investment of time, there are a number of advantages over mRNA injections. Namely, conditional expression can be achieved at any age using a range of available induction systems. Our current heat shock promoter-based approach offers researchers the opportunity to restrict expression to specific time points during development, which should enable the study of dynamic changes in proximal interactome over the course of early development. While we have not explicitly demonstrated this here, crosses with other lines can be performed to investigate proximity interactions in diverse genetic backgrounds. Additional transgenic systems could also be generated that enable tissue-specific expression by the use of tissue-specific promoters and enhancers, though this may require additional optimization (e.g. to enrich the tissues/cells labeled in order to improve the signal-to-noise ratio). The studies of Pronobis *et al*. (26) have demonstrated that biotin ligase-based proximity labeling is also feasible in adult zebrafish, and our heat shock promoter transgenics system should be appropriate to study proximal interactions in adults. Whether the biotin will need to be supplied through intraperitoneal injection as in Pronobis *et al*., or whether there will be sufficient biotin in the water to enable biotinylation by the more active TurboID enzyme will however need to be defined.

While we demonstrate that both the TurboID and miniTurbo enzymes can be used for performing BioID in the zebrafish, there were some differences. We found that in transgenic fish lines the miniTurbo fusion constructs showed higher expression as compared to TurboID constructs. However, levels of biotinylation as well as protein recovery was higher using TurboID. The expression levels combined with the protein recovery suggests that while the smaller size of miniTurbo may result in better expression, the higher biotinylation efficiency of TurboID compensates for the lower expression. These factors should be considered when deciding which enzyme to use. When using short labeling times, such as when using inducible expression to query different timepoints during development, the higher efficiency of TurboID should allow for shorter labeling periods without significant decreases in sensitivity. On the other hand, if performing non-inducible tissue specific labelling the higher expression levels of miniTurbo may provide equally useful results. Finally, others have found that the high efficiency of TurboID leads to toxicity when expressed constitutively and ubiquitously in flies, presumably through scavenging free biotin so efficiently (9). While we have not found this to be the case in our zebrafish experiments so far, we cannot rule it out, and it is possible that miniTurbo may be more appropriate in these cases.

In conclusion, we have developed and benchmarked tools for reproducible proximity-dependent biotinylation in zebrafish, that should be of general use to expand BioID in this important model vertebrate system. To allow the zebrafish community to easily take advantage of this system, we make available the pCS2+ mRNA expression vectors containing the TurboID and miniTurbo sequences for easy generation of mRNA coding for bait fusion proteins without the time consuming need to develop transgenic animals. We also make available Tol2Kit amenable vectors coding for TurboID and miniTurbo to enable transgenic expression of tagged proteins of interest (Supplementary Figure 6).

## Abbreviations

AMP: Adenosine monophosphate
APEX: enhanced ascorbate peroxidase
BASU: Bacillus subtilis biotin ligase
BFDR: Bayesian false discovery rate
BioID: proximity-dependent biotin identification
FDR: False discovery rate
GFP: Green fluorescent protein
GO: Gene ontology
HPF: hours post fertilization
LMNA: lamin A
MS: Mass spectrometry
WT: Wild type

## Acknowledgements

We thank Brett Larsen and Cassandra Wong for assistance with mass spectrometry and Payman Samavarchi-Tehrani for helpful discussion and technical advice. We also thank the SickKids Zebrafish Facility staff, Alejandro Salazar, Elyjah Schimmens, Melissa Westaway, and Scott Knox for animal husbandry. Work in the Gingras lab was supported by a Canadian Institutes of Health Research (CIHR) Foundation Grant (FDN 143301). Proteomics work was performed at the Network Biology Collaborative Centre at the Lunenfeld-Tanenbaum Research Institute, a facility supported by Canada Foundation for Innovation funding, by the Ontario Government, and by Genome Canada and Ontario Genomics (OGI-139). Work in the Scott lab was supported by a Canadian Institutes of Health Research (CIHR) project Grant (PJT 153000) and a Natural Sciences and Engineering Research Council (NSERC) Discovery Grant (RGPIN-2017-06502). S.M.R. is supported by a SickKids Restracomp scholarship and T.C.B. was supported by Dow Graduate Research and Lester Wolfe Fellowships. A.-C.G. is the Canada Research Chair in Functional Proteomics and the Lea Reichmann Chair in Cancer Proteomics.

## Data availability

Datasets consisting of raw files and associated peak lists and results files have been deposited in ProteomeXchange through partner MassIVE (http://proteomics.ucsd.edu/ProteoSAFe/datasets.jsp) as complete submissions. Additional files include the sample description, the peptide/protein evidence and the complete SAINTexpress output for each dataset, as well as a “README” file that describes the dataset composition and the experimental procedures associated with each submission. The different datasets generated here were submitted as independent entries. All submissions are password-protected until publication. Password: zebrafish.

MassIVE deposition 1: Rosenthal_Zebrafish_TurboID_P88_SAINT5425_TurboID_injections_2021 This dataset consists of 14 raw MS files and associated peak lists and result files comprising of mRNA injections of the TurboID fused baits.

ftp://MSV000087425@massive.ucsd.edu

*PXD026007*

MassIVE deposition 2: Rosenthal_Zebrafish_TurboID_P88_SAINT5427_miniTurbo_injections_2021 This dataset consists of 13 raw MS files and associated peak lists and result files comprising of mRNA injections of the miniTurbo fused baits.

ftp://MSV000087428@massive.ucsd.edu

*PXD026021*

MassIVE deposition 3: Rosenthal_Zebrafish_TurboID_P88_SAINT5428_TurboID_transgenics_2021 This dataset consists of 9 raw MS files and associated peak lists and result files comprising of transgenics (heat-shock) analysis of the TurboID fused baits.

ftp://MSV000087429@massive.ucsd.edu

*PXD026022*

MassIVE deposition 4:

Rosenthal_Zebrafish_TurboID_P88_SAINT5429_miniTurbo_transgenics_2021

This dataset consists of 9 raw MS files and associated peak lists and result files comprising of transgenics (heat-shock) analysis of the miniTurbo fused baits.

ftp://MSV000087430@massive.ucsd.edu

*PXD026023*

MassIVE deposition 5:

Rosenthal_Zebrafish_TurboID_P88_SAINT5454_TurboID_transgenics_conditions_2021 This dataset consists of 22 raw MS files and associated peak lists and result files comprising of transgenics (heat-shock) analysis of the TurboID and comparison of conditions (biotin and heat shock timing and concentration).

ftp://MSV000087431@massive.ucsd.edu

*PXD026024*

## Supplemental Figures

**Supplementary Figure 1.**
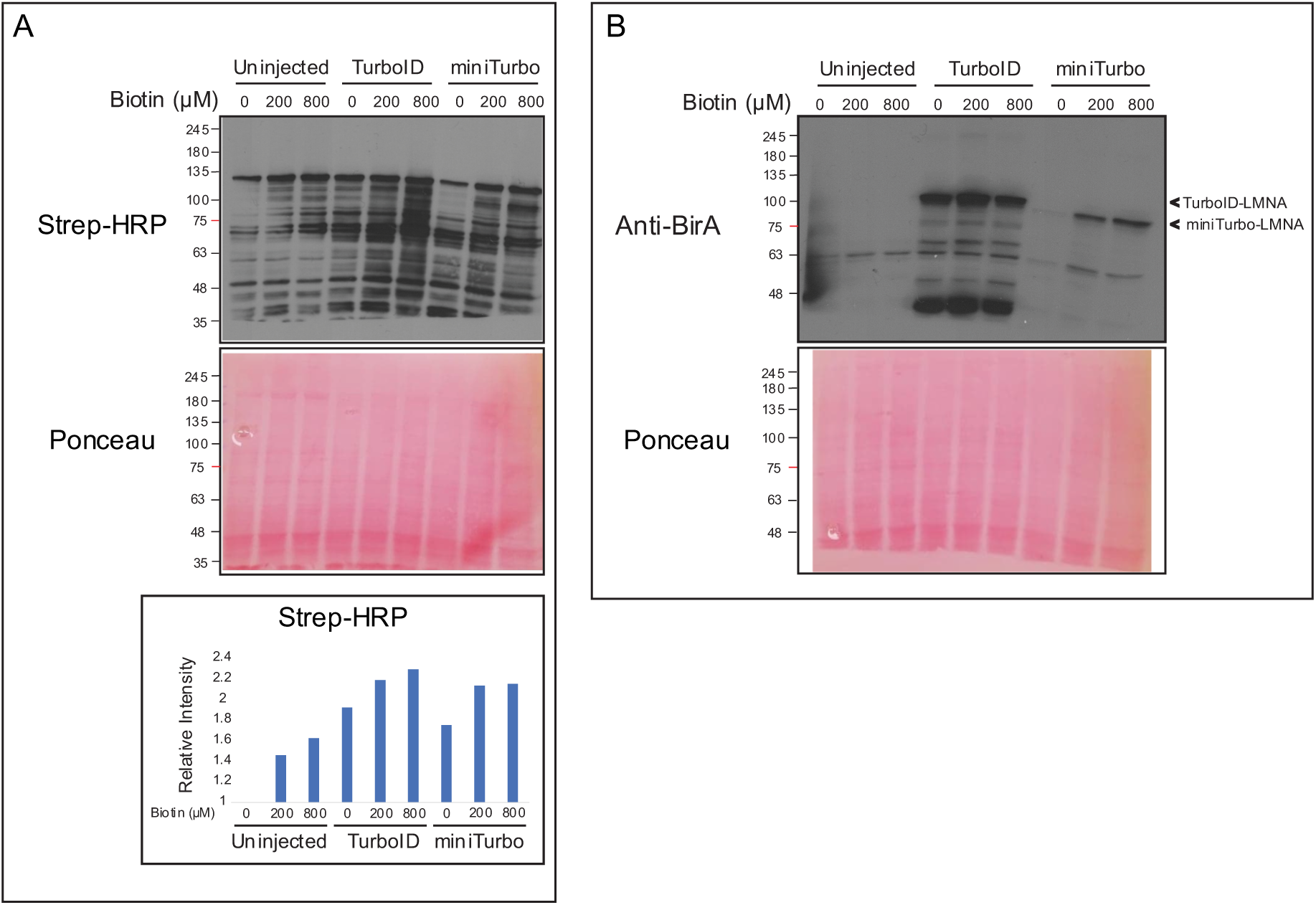
Western Blot analysis shows that mRNA injection-based expression of biotinylating enzymes produces increased protein biotinylation as biotin concentration is increased. (A) Streptavidin-HRP was used to probe proteins from pooled whole embryo lysates of zebrafish embryos injected with TurboID-LMNA, miniTurbo-LMNA, and control uninjected embryos and incubated with various biotin supplementation concentrations for 48 hours. (B) An anti-BirA antibody was used to probe for TurboID and miniTurbo.

**Supplementary Figure 2.**
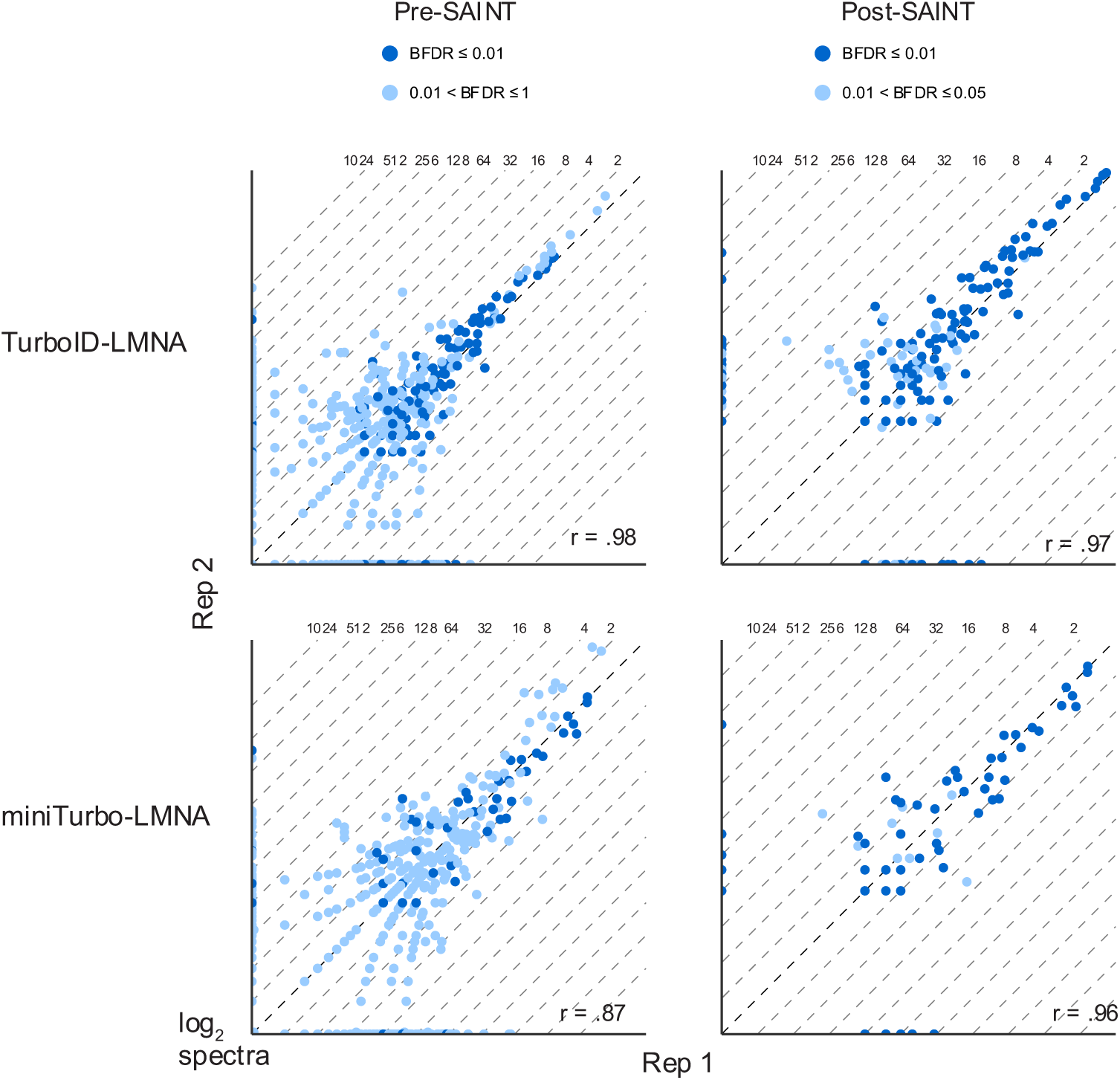
Proteins identified at 12 hpf using mRNA injection based BioID experiments show high correlation between biological replicates. BioID experiments using mRNA injections of TurboID-LMNA and miniTurbo-LMNA were performed in biological duplicates. Injected embryos were incubated with 800 µM biotin for 12 hours. Correlation Pre-SAINT shows all proteins identified with ≥ 2 unique peptides and an iProphet probability of ≥ 95%. Post-SAINT correlation includes the same parameters as for pre-SAINT with an additional requirement for a BFDR ≤ 5%.

**Supplementary Figure 3.**
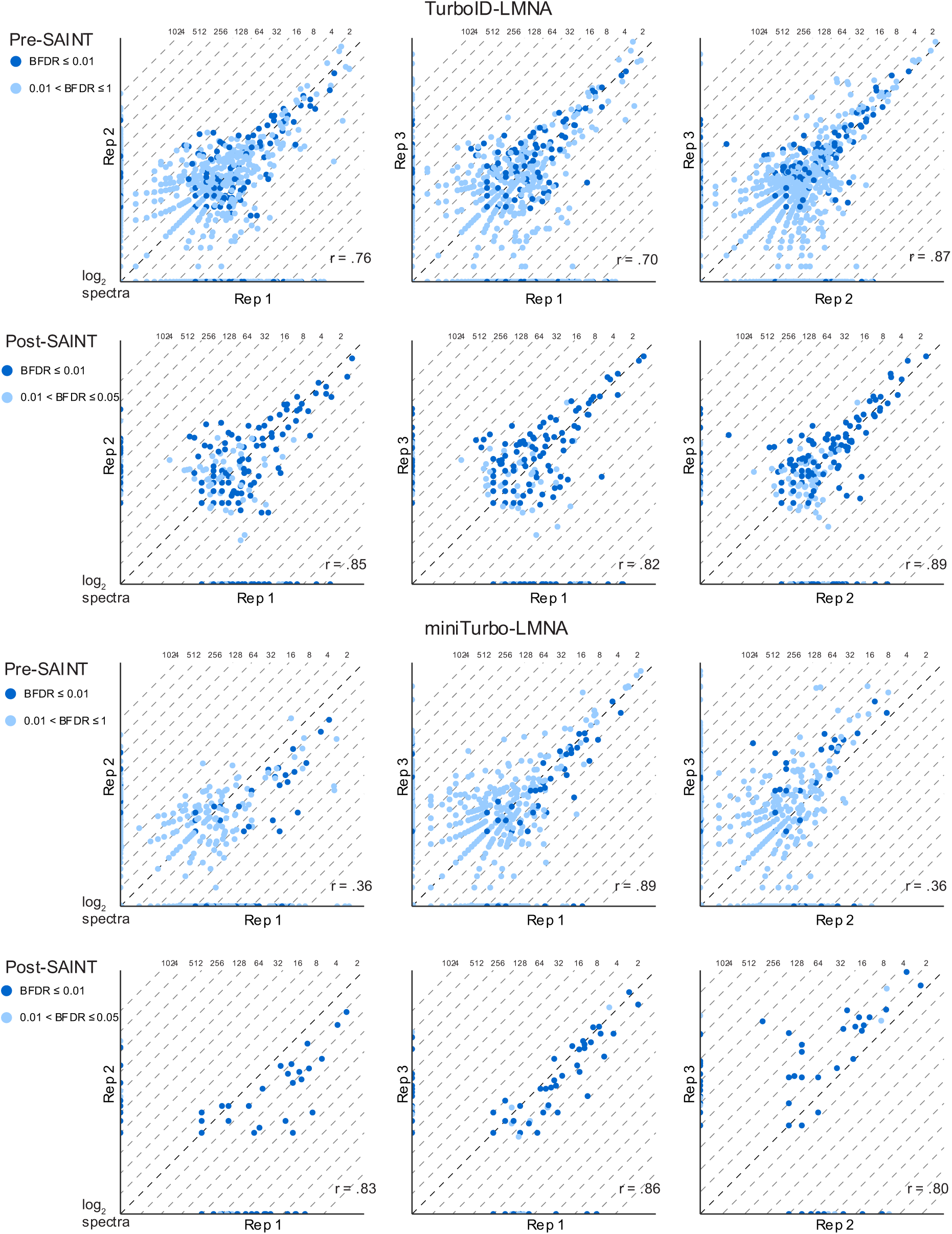
Proteins identified at 48 hpf using mRNA injection based BioID experiments show varying degrees of correlation between biological replicates. BioID experiments using mRNA injections of TurboID-LMNA and miniTurbo-LMNA were performed in biological triplicates. Injected embryos were incubated with 800 µM biotin for 48 hours. Correlation Pre-SAINT shows all proteins identified with ≥ 2 unique peptides and an iProphet probability of ≥ 95%. Post-SAINT correlation includes the same parameters as for pre-SAINT with an additional requirement for a BFDR ≤ 5%.

**Supplementary Figure 4.**
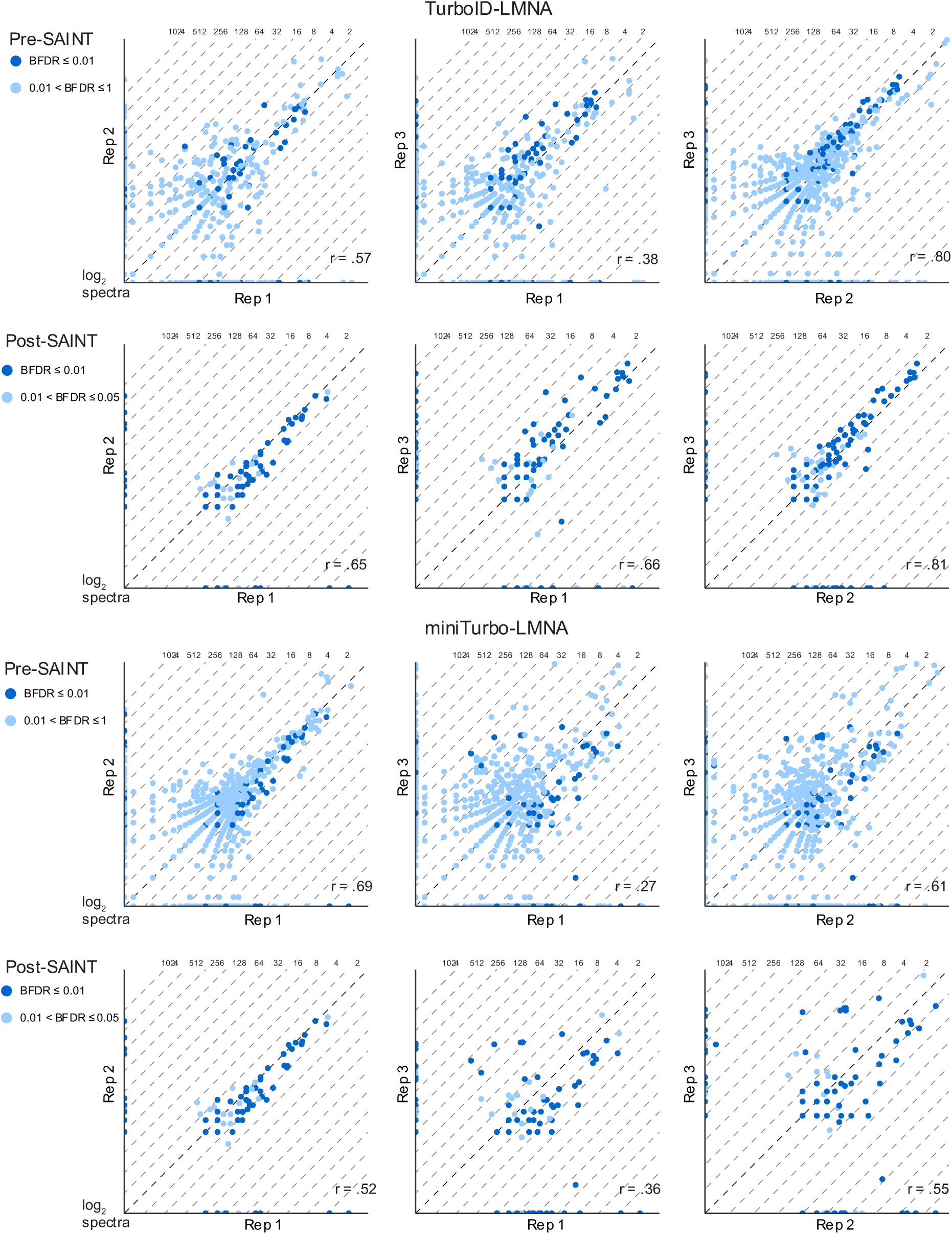
Proteins identified at 72 hpf using heat shock inducible transgenic based BioID experiments show a lower degree of correlation between biological replicates than mRNA injection based experiments. BioID experiments using heat shock induction of TurboID-LMNA and miniTurbo-LMNA at 60 hpf for one hour were performed in biological triplicates. Induced embryos were incubated with 800 µM biotin for from 0 hpf to collection at 72 hpf. Correlation Pre-SAINT shows all proteins identified with ≥ 2 unique peptides and an iProphet probability of ≥ 95%. Post-SAINT correlation includes the same parameters as for pre-SAINT with an additional requirement for a BFDR ≤ 5%.

**Supplementary Figure 5.**
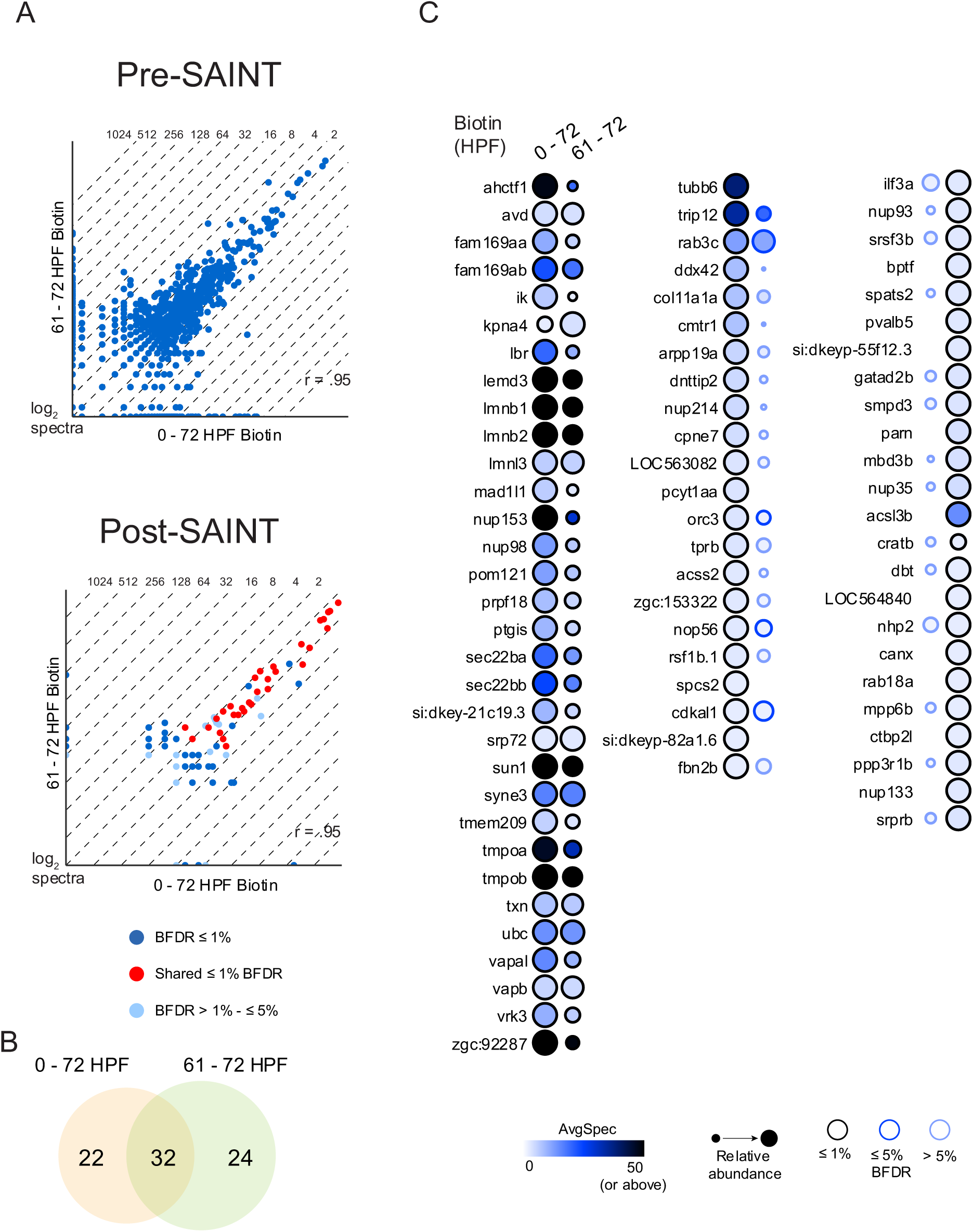
The timing of biotin supplementation has a small influence on the recovery of proximal partners in transgenic experiments. Using the transgenic TurboID-LMNA line, we investigated the effects of changes in biotin supplementation time on protein identifications. BioID experiments using heat shock induction of TurboID-LMNA at 60 hpf for one hour were performed in biological duplicates. Induced embryos were incubated with 800 µM biotin for from 0 hpf or 61 hpf to collection at 72 hpf. (A) Correlation Pre-SAINT shows all proteins identified with ≥ 2 unique peptides and an iProphet probability of ≥ 95%. Post-SAINT correlation includes the same parameters as for pre-SAINT with an additional requirement for a BFDR ≤ 5%. (B) Venn diagram, showing the overlap between proteins identified with a BFDR ≤ 1% in the two conditions. (C) Dotplot showing all the proteins identified with a BFDR ≤ 1%.

**Supplementary Figure 6.**
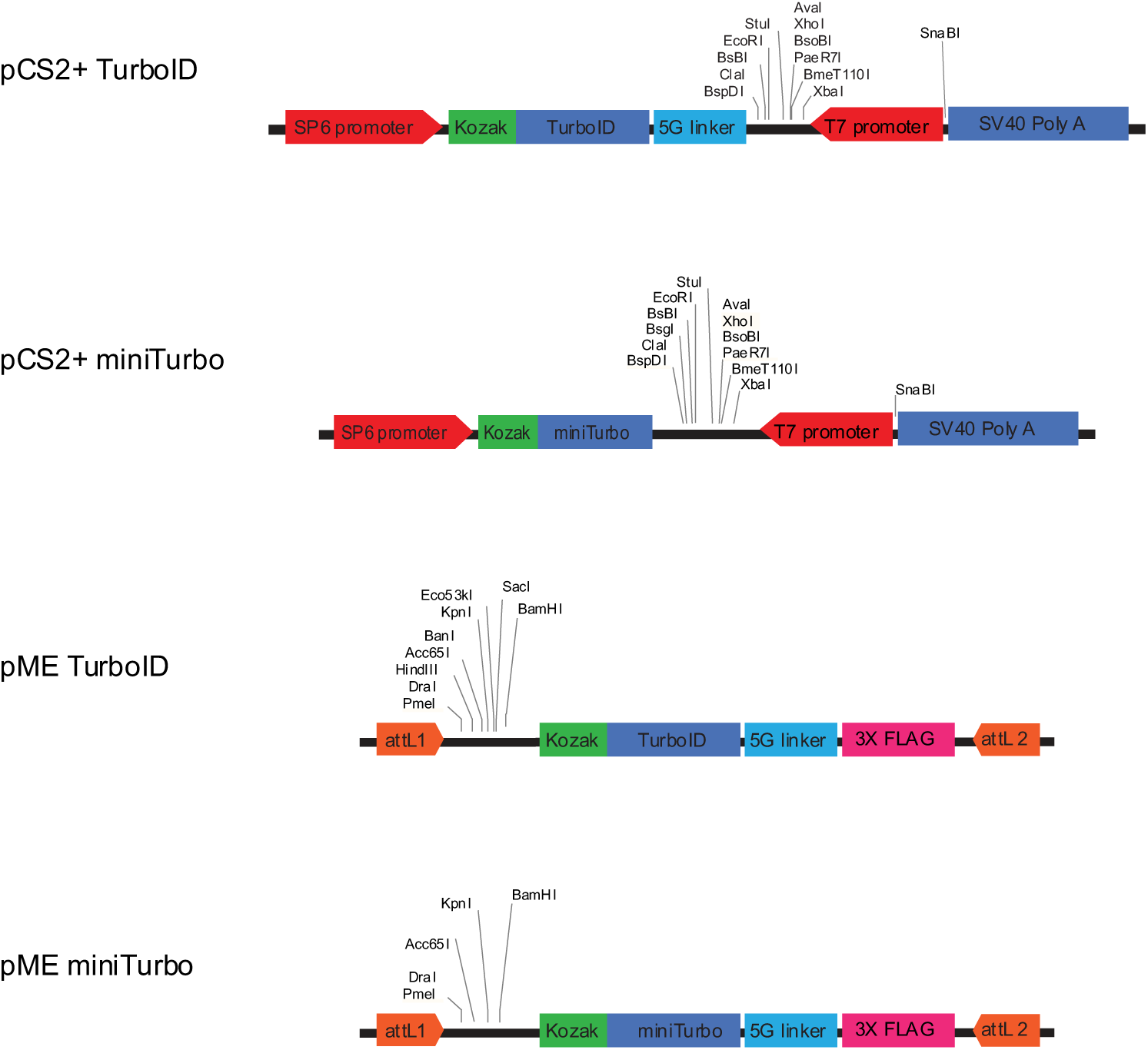
Diagram of cassettes used in vectors available for performing mRNA injection or transgenic based BioID in zebrafish. To make the TurboID and miniTurbo constructs used in this paper available to the community, we cloned these constructs into the pCS2+ and pME vectors. The pCS2+ vector can be used for *in vitro* transcription of capped mRNA. This vector contains the TurboID or miniTurbo enzyme downstream of a SP6 promoter site and with a zebrafish consensus Kozak sequence at the start site. Downstream of the enzyme sequence is a multiple cloning site for insertion of a gene of interest, this is preceded by a linker in the TurboID construct. Both constructs contain a T7 promoter site and SV40 Poly A sequence at the 3’ end of the cassette. For generation of transgenic zebrafish, we provide TurboID and miniTurbo cassette within the Gateway cloning based Tol2 Kit pME middle entry vector. These vectors contain the Kozak and enzyme sequence as in the pCS2+ vectors with the addition of a linker and 3X FLAG at the 3’ end of the enzyme. The cassette is flanked by Gateway attL sites.

## Supplementary Table Descriptions

**Supplementary Table 1. SAINT analysis output tables for mRNA injection based BioID experiments.** The SAINT analysis output for the four experiments (TurboID-LMNA 12 hour labeling, TurboID-LMNA 48 hour labeling, miniTurbo-LMNA 12 hour labeling, and miniTurbo-LMNA 48 hour labeling) performed using the mRNA injection-based method. The analysis was performed using the SAINTexpress integration on ProHits.

**Supplementary Table 2. SAINT analysis output tables for inducible transgenic based BioID experiments.** The SAINT analysis output for the two experiments (TurboID-LMNA and miniTurbo-LMNA) performed using the heat shock induction-based method. The analysis was performed using the SAINTexpress integration on ProHits.

**Supplementary Table 3. Human orthologs of the identified high-confidence interactors.** The zebrafish genes associated with the proteins identified with a BFDR ≤ 1% were searched for human orthologs. The zebrafish genes and the identified human orthologs are listed.

**Supplementary Table 4. Gene Ontology (GO) enrichment analysis results.** The human orthologs of the identified high-confidence interactors were used to perform GO enrichment analysis. The GO analysis data source used GO Cellular Component (CC) Terms containing 5–500 genes.

**Supplementary Table 5. Comparison of high-confidence interactors from this study with previous datasets.** We compared the human orthologs of the zebrafish high-confidence interactors with the high confidence interactors from previously published studies. The high-confidence interactors from all datasets were searched for genes annotated with the GO CC terms nuclear membrane and nuclear envelope.

**Supplementary Table 6. Source data for the summed spectral counts comparison.** To compare the identification of the high-confidence interactors identified using transgenic TurboID-LMNA with 800 µM biotin supplementation form 0 hpf with experiments using different biotin supplementation times, concentrations, and transgene induction parameters, we summed the spectra identified for these 54 proteins across two replicates.

**Supplementary Table 7. SAINT analysis output tables for inducible transgenic TurboID-LMNA biotin supplementation time, concentration, and transgene induction parameter comparison experiments.** The SAINT analysis output for the experiment using different biotin supplementation times, concentrations, and transgene induction parameters. The analysis was performed using the SAINTexpress integration on ProHits.

**Supplementary Table 8. GO enrichment analysis for experiment comparing biotin supplementation at 0 hpf versus 61 hpf.** GO enrichment analysis was performed using as input the human orthologs of the high-confidence interactors identified in transgenic TurboID-LMNA BioID experiments using biotin supplementation ‘pre-load’ from 0 hfp or only from 61 hpf. The GO analysis data source used GO Cellular Component (CC) Terms containing 5–500 genes.

